# CD40 agonistic-monovalent streptavidin fusion antibody for targeted neoantigen peptide delivery and potent cancer vaccination

**DOI:** 10.1101/2025.07.30.663779

**Authors:** Dahee Jung, Xiaoying Cai, Ziye Wan, Nawon Lee, Neal T Ramseier, Ying Hu, Seung-Oe Lim, Steve Seung-Young Lee

## Abstract

Cancer vaccines targeting patient-derived neoantigens offer great promise for personalized cancer therapy but face challenges in achieving targeted delivery to antigen-presenting cells (APCs) to elicit robust and durable cancer-specific immune responses. We synthesized an anti-mouse CD40 agonistic-monovalent streptavidin fusion antibody (αCD40-mSAs), which enables targeted delivery of biotinylated neoantigen peptides to APCs in draining lymph nodes (dLNs). We confirmed mSA expression on the engineered antibody and its strong binding affinities to mouse CD40 and biotin. Advanced microscopy demonstrated that αCD40-mSAs enhances homing to dLNs and intracellular delivery of neoantigen peptides to critical APC subsets, such as cDC1. The potent agonistic effects of αCD40-mSAs on dendritic cell maturation, activation, and antigen presentation were verified through *in vitro* assays. Vaccination with αCD40-mSAs elicited robust cancer-specific CD8⁺ T cell responses, leading to significant tumor regression and prevention in a mouse tumor model. These results support αCD40-mSAs as an ‘all-in-one’ vaccine delivery platform with multifunctional immunopharmacological advantages and strong translational potential for personalized cancer vaccination.

**Teaser:** αCD40-mSAs is an engineered anti-CD40 agonistic antibody designed to enhance cancer vaccine delivery.

## Introduction

Cancer vaccines targeting neoantigens derived from individual patients’ mutanomes represent a rational and promising strategy for personalized cancer immunotherapy (*1*, *2*). Unlike vaccines directed at overexpressed or tumor-associated antigens, neoantigens are not subject to central or peripheral immune tolerance, making them ideal targets for eliciting cancer-specific immune responses while minimizing the risk of adverse effects such as autoimmunity (*1*, *3*, *4*).

Recent advances in peptide and mRNA synthesis have enabled their widespread use as antigenic materials for cancer vaccines, offering rapid customization and cost-effective production for clinical applications (*5*, *6*). However, naked mRNA is unsuitable for direct administration due to its low stability and biosafety concerns. As a result, mRNA-based vaccines require encapsulation in nanoparticle carriers such as liposomes, which presents regulatory and manufacturing challenges when formulating personalized vaccines (*7*). In contrast, synthetic long peptides (SLPs) exhibit higher *in vivo* stability and can be directly administered with adjuvants following simple mixing, making them more suitable for personalized cancer vaccines in clinical trials (*8*, *9*). Several clinical studies using neoantigenic peptides have demonstrated some clinical benefit, including the induction of anti-tumor immune responses and long-term prevention of tumor relapse, with a median follow-up of 40.2 months (*10*). However, only a subset of patients benefits from such vaccines, as seen in many other peptide vaccine trials (*11*, *12*). Although some patients exhibit noticeable responses to cancer vaccines, most cases fail to establish robust and durable cancer-specific CD8⁺ T cell immunity (*10*, *11*), which is critical for achieving effective anti-tumor vaccination outcomes (*13*).

Subcutaneously (s.c.) or intramuscularly (i.m.) administered unformulated peptides and adjuvants are generally taken up by local antigen-presenting cells (APCs), including dermal and epidermal dendritic cells (DCs) and macrophages (*14*, *15*). These APCs then migrate to the draining lymph nodes (dLNs) to initiate T cell priming (*14*). However, this non-specific, passive delivery pathway often results in poor immunopharmacological efficacy, including inefficient cellular uptake, limited transport to dLNs, suboptimal antigen presentation, and inconsistent immune activation. These limitations can impair vaccine efficacy and increase the risk of off-target immune responses (*16*). To overcome these limitations, targeted delivery of vaccines to APCs in dLNs has been actively investigated (*17*, *18*). Notably, small molecules like naked peptides (<40 kDa) primarily enter the bloodstream after s.c. injection, whereas larger molecules such as IgG antibodies (>84 kDa) preferentially drain into the lymphatic system (*19–21*). Furthermore, simple mixtures of peptides and adjuvants often fail to co-deliver into the same APCs due to differences in their biophysical properties and pharmacokinetics, limiting effective APC activation and antigen presentation (*22*, *23*).

To address this pharmacological challenge of current peptide vaccine approaches, various nanoparticle formulations have been developed for the co-delivery of peptides and adjuvants. (*24–27*). However, such platforms face translational barriers in personalized vaccine settings, particularly in manufacturing and regulatory compliance of patient-tailored formulations. Antibody-based peptide vaccine strategies have also been explored, for instance, DC-targeting antibodies such as anti-DEC-205 and CLEC9A engineered to express tumor-associated antigens (e.g., NY-ESO-1) *via* protein–antibody recombination (*28–31*). However, these approaches often involve complex synthesis and require the co-administration of adjuvants, rendering them suboptimal for the rapid and customizable manufacturing needed for effective patient-specific vaccination. Thus, there remains an urgent need for an “All-in-One” delivery platform capable of: (1) targeting APCs (e.g. DCs (*32*, *33*)) in dLNs, (2) enabling efficient cellular internalization in APCs, and (3) co-delivering antigen and adjuvant in a single formulation. Such a platform must also maintain flexibility to accommodate diverse neoantigen peptides, enabling clinical translation of personalized cancer vaccination.

CD40 is a TNF receptor superfamily member expressed on the plasma membrane of various APCs, including DCs and macrophages in dLNs (*34–36*). As a key co-stimulatory molecule involved in both innate and adaptive immune responses, CD40 activation using an anti-CD40 agonistic antibody (αCD40) promotes DC maturation and T cell priming in dLNs, leading to robust anti-tumor immunity (*34*, *37*, *38*). Moreover, αCD40 bound to membrane-expressed CD40 is efficiently internalized into early endosomes of DCs. Additionally, IgG antibodies, including αCD40, exhibit natural dLN-homing properties, with rapid lymphatic drainage following s.c. injection (*39*), further supporting the use of αCD40 as a dLN-homing vaccine carrier.

In this study, we leveraged the immunopharmacological advantages of αCD40, including its dLN homing, APC targeting, cellular internalization, and immunostimulatory capacity, as a delivery vehicle for cancer neoantigen peptides. To enable flexible peptide conjugation, we engineered αCD40 to express monovalent streptavidins (αCD40-mSAs) at the C-terminus of both light chains. This construct allows for stable, high-affinity binding of biotinylated neoantigen peptides *via* the mSA-biotin interaction. Advanced biotinylation techniques allow site-specific labeling of peptides with biotin (*40*) without altering their amino acid sequence or biophysical characteristics (*41*). Therefore, αCD40-mSAs represents a “plug & play” delivery system for personalized cancer vaccination. We demonstrate that αCD40-mSAs enhances peptide delivery to APCs in dLNs and promotes the induction of neoantigen-specific T cell responses, resulting in both therapeutic and preventive anti-tumor effects in mouse tumor models. These findings support αCD40-mSAs as a promising neoantigen peptide delivery platform for personalized cancer vaccination.

## Results

### Synthesis and characterization of engineered anti-CD40 agonistic-monovalent streptavidin fusion antibody (αCD40-mSAs)

We designed anti-CD40 agonistic antibody to express monovalent streptavidins at the C-terminus of light chains with a short peptide space, referred to as αCD40-mSAs (**Fig. 1A**). Using amino acid sequences from the complementarity-determining regions (CDRs) of the anti-mouse CD40 agonistic antibody (clone FGK4.5) and from monovalent streptavidin (mSA) (*42*), we reconstructed cloning vectors based on mouse IgG2a, which shares similar Fcγ receptor (FcγR)-binding properties with human IgG1 (*43*, *44*). Recombinant αCD40 and αCD40-mSAs were produced using a CHO cell expression system, purified using a FPLC system, yielding over 50 mg/L and 100 mg/L, respectively. After exchanging buffer to PBS and confirming a low endotoxin level (<0.5 EU), the antibodies were aliquoted into individual vials and stored at -80 °C.

**Figure 1.**
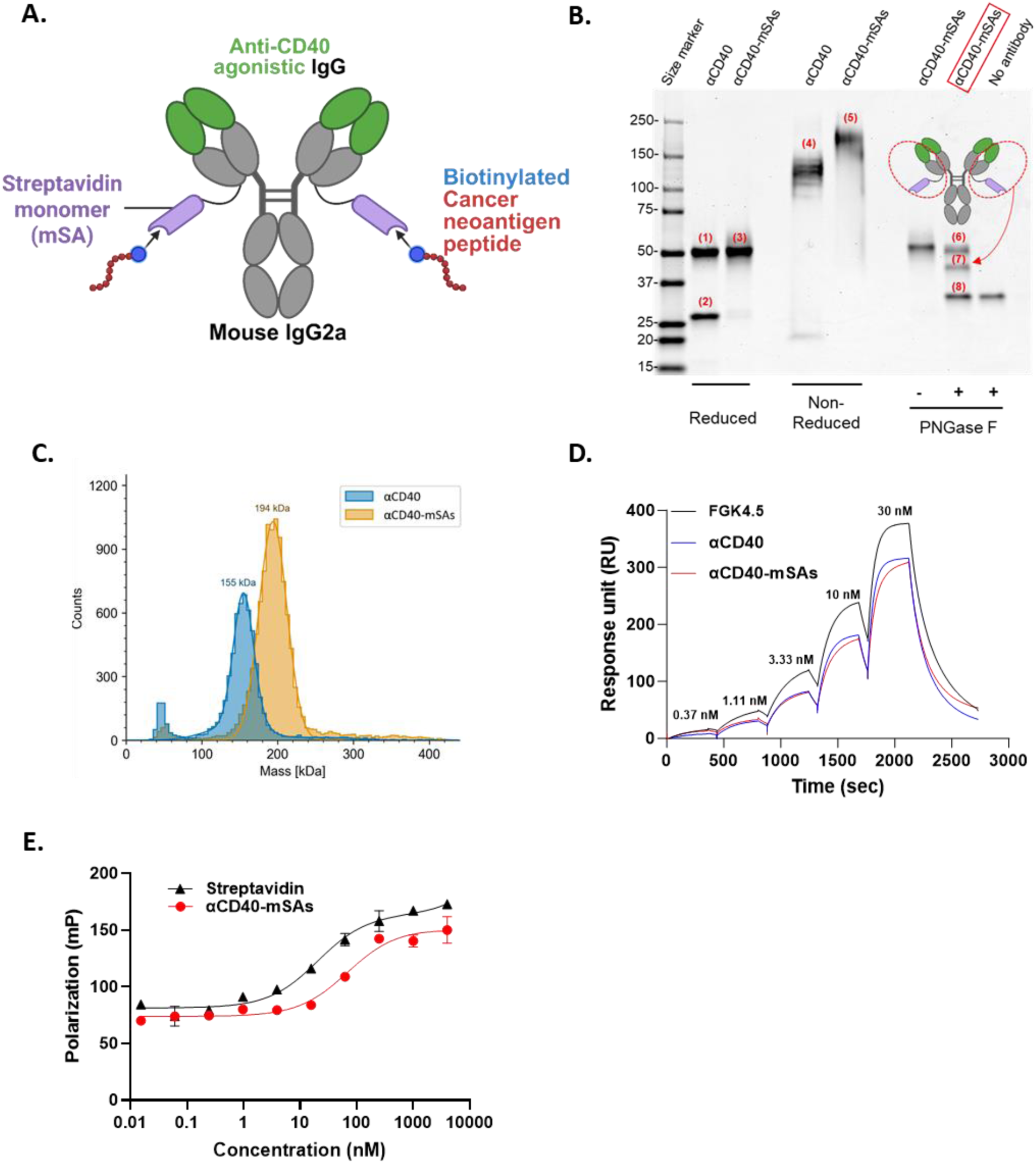
Design and biophysical characterization of αCD40-mSAs. (A) Schematic illustration of biotinylated neoantigen peptide-loaded αCD40-mSAs. (B) SDS-PAGE analysis. Under reducing conditions: (1) heavy and (2) light chains of αCD40; (3) heavy and light chains of αCD40-mSAs. Under non-reducing conditions: (4) αCD40 and (5) αCD40-mSAs. Under reducing conditions after deglycosylation: (6) heavy and (7) light chains of deglycosylated αCD40-mSAs; (8) PNGase F. (C) Mass photometry (MP) measurement of recombinant αCD40 and αCD40-mSAs, showing peak molecular weights of ∼155 and 194 kDa, respectively. (D) Surface plasmon resonance (SPR) sensorgram showing high binding affinity of recombinant αCD40 and αCD40-mSAs to mouse CD40, comparable to that of commercial αCD40 (clone FGK4.5). (E) Fluorescence polarization (FP) of biotin-FITC in the presence of varying concentrations of streptavidin or αCD40-mSAs, demonstrating biotin-binding functionality.

The αCD40-mSAs construct enables the loading of biotinylated neoantigen peptides specifically at the C-termini of its light chains *via* the mSA-biotin interaction, while preserving the variable and Fc regions for binding to CD40 and Fcγ receptors (FcγRs), respectively. The agonistic effect of αCD40 on APCs is primarily mediated through recognition of its Fc domain by FcγRs on other immune cells, which induces CD40 clustering on APCs and promotes their activation and maturation (*34*). Therefore, preserving the Fc domain in αCD40 is critical for inducing its agonistic effect on APC activation as a molecular adjuvant. SDS-PAGE analysis under reducing conditions showed distinct bands for the heavy chain (∼50 kDa, labeled (1) in **Fig. 1B**) and light chain (∼25 kDa, (2)) of αCD40. In contrast, αCD40-mSAs showed a single band (at ∼50 kDa (3)) due to glycosylation of mSAs on the light chains. Under non-reducing conditions, αCD40-mSAs exhibited an increased molecular weight (∼200 kDa, (5)) compared to the parental αCD40 (∼150 kDa, (4)). After deglycosylation with PNGase F and reduction, αCD40-mSAs yielded bands for the heavy chain (∼50 kDa, (6)), the light chain (∼40 kDa, (7)), and the PNGase F enzyme (∼30 kDa, (8)). The increase in the light chain molecular weight from ∼25 kDa (2) to ∼40 kDa (7) indicates successful mSA expression. Molecular weight profiles of αCD40 (peak: 155 kDa) and αCD40-mSAs (peak: 194 kDa) were further confirmed by mass photometry (MP) (**Fig. 1C**). Surface plasmon resonance (SPR) analysis demonstrated high-affinity binding of both αCD40 and αCD40-mSAs to mouse CD40, as shown in representative sensorgrams (**Fig. 1D**). The equilibrium dissociation constants (K_d_) for the commercial αCD40 agonist antibody (clone FGK4.5), the synthesized αCD40, and αCD40-mSAs were comparable, measured at 8.4, 12.3, and 12.7 nM, respectively (**Table 1**). Additional ELISA assays also confirmed the interaction with mouse CD40 (**Fig S1. A, B**). A fluorescence polarization (FP) assay was used to evaluate the interaction between αCD40-mSAs and biotin. Gradual addition of αCD40-mSAs to a biotin-FITC solution reduced the Brownian motion of free biotin-FITC due to strong mSA-biotin interactions, leading to increased FP signal (**Fig. 1E**). The determined K_d_ values were 69.23 nM for αCD40-mSAs and 20.56 nM for recombinant streptavidin. An ELISA assay further validated the binding of biotin to αCD40-mSAs (**Fig S1. C**). These biophysical characterizations of αCD40-mSAs confirm its successful synthesis, biotin-binding functionality, and CD40-target specificity.

**Table 1.** Kinetic constants for binding to mouse CD40.

|  | ka (1/Ms) | kd (1/s) | K<sub>d</sub> (nM) |
| --- | --- | --- | --- |
| FGK 4.5 | 7.8E+05 (± 1.8E+04) | 6.5E-03 (± 7.9E-05) | 8.4 (± 0.13) |
| αCD40 | 1.0E+06 (± 2.5E+04) | 1.3E-02 (± 2.2E-04) | 12.3 (± 0.13) |
| αCD40-mSAs | 2.5E+06 (± 2.8E+05) | 3.2E-02 (± 3.4E-03) | 12.7 (± 0.27) |

\*ka: Association constant, kd: Dissociation constant, and K<sub>d</sub>: Equilibrium dissociation constant

### Induction of *in vivo* antigen-specific CD8⁺ T cell responses by biotinylated peptide-loaded αCD40-mSAs

We first evaluated whether αCD40-mSAs could elicit a potent antigen-specific CD8⁺ T cell response, a critical effector for therapeutic vaccination. A biotinylated SLP containing the OVA257-264 epitope (SIINFEKL) (Bio-OVA; detailed sequence provided in **Table S1**) was used as a model neoantigen. C57BL/6 mice were vaccinated with different Bio-OVA formulations *via* footpad injection according to the schedule shown in **Fig. 2A**. Seven days post-injection, vaccinated mice were sacrificed and dLNs and spleens were harvested. To identify activated, OVA-specific cytotoxic T cells, we performed flow cytometry using PE-labeled H2-Kb/OVA (SIINFEKL) MHC class I tetramer along with anti-CD44 and anti-CD8 antibodies. The group vaccinated with αCD40-mSAs-Bio-OVA exhibited a significantly higher percentage of OVA tetramer⁺ CD44⁺ CD8⁺ T cells in the both dLNs and spleens compared to other vaccine groups (**Fig. 2B**). This adaptive immune response was associated with a marked increase in the absolute number of OVA-specific cytotoxic CD8⁺ T cells following a single dose of αCD40-mSAs-Bio-OVA (**Fig. 2C**). Representative flow cytometry plots illustrate the extent of cytotoxic T cell expansion in response to the OVA neoantigen across different vaccine groups (**Fig. 2D**). These results suggest that αCD40-mSAs enhance the immunopharmacological properties of neoantigen peptides, leading to the induction of robust, neoantigen-specific CD8⁺ T cell immunity.

**Figure 2.**
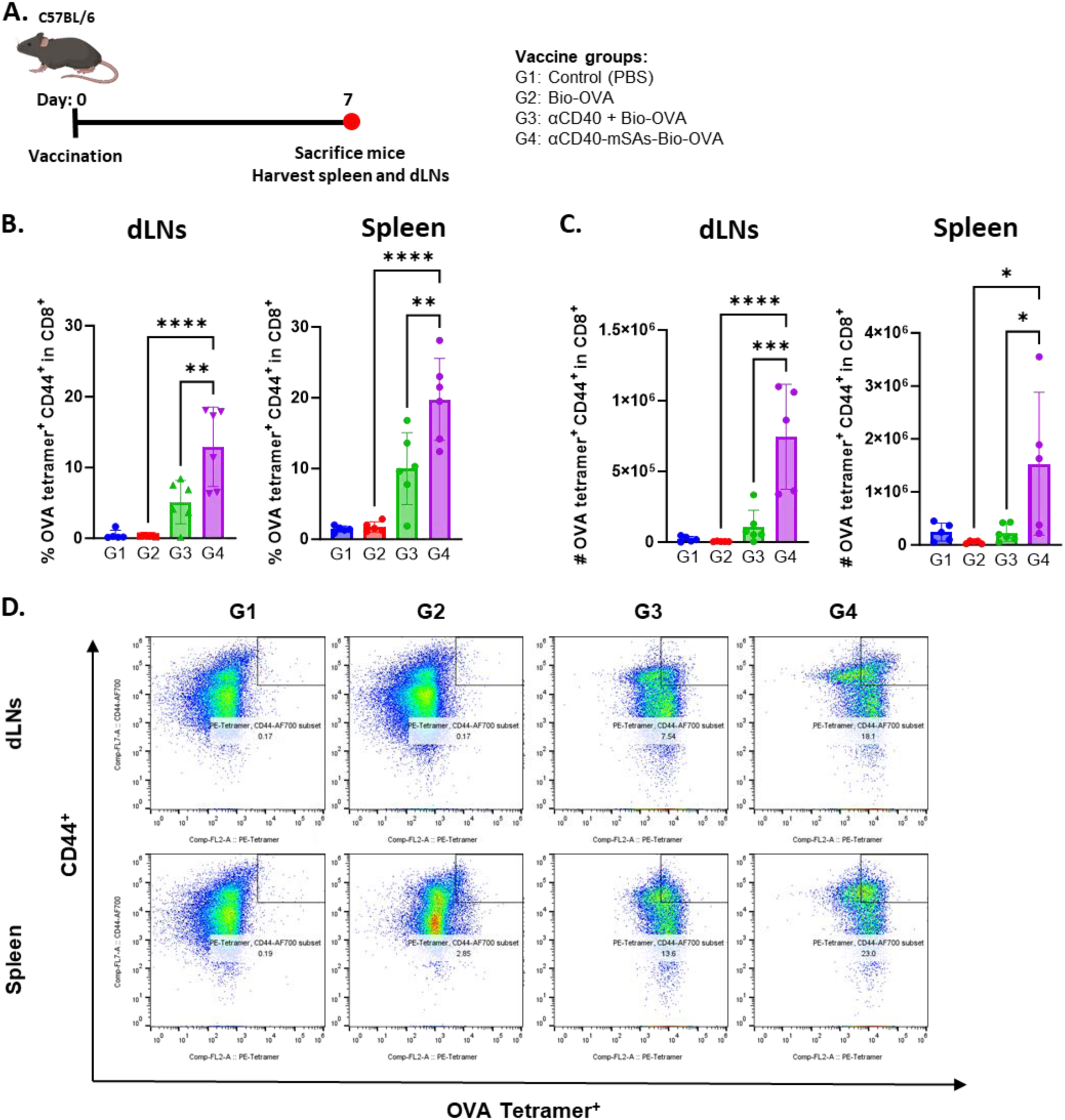
Induction of neoantigen-specific CD8⁺ T cell responses by a single vaccination with αCD40-mSAs *in vivo*. (A) Experimental timeline for single vaccination and tissue collection. (B) Frequency and (C) absolute number of OVA tetramer⁺ CD44⁺ CD8⁺ T cells in the draining lymph nodes (dLNs) and spleen. (D) Representative flow cytometry plots of OVA tetramer⁺ CD44⁺ CD8⁺ T cells in the dLNs (top) and spleen (bottom). \**p* < 0.05; \*\**p* < 0.01; \*\*\**p* < 0.001; \*\*\*\**p* < 0.0001.

### dLN-homing and APC-targeted delivery of neonatigen peptide by αCD40-mSAs

To evaluate the neoantigen delivery efficacy of the vaccine formulations studied above, we conducted an *in vivo* biodistribution study using fluorophore-labeled Bio-OVA and αCD40 or αCD40-mSAs. Mice were injected with fluorescent vaccine formulations *via* footpad, and the popliteal (PO) dLNs were harvested 90 minutes post-injection. Confocal fluorescence microscopy was used to detect signals from injected Bio-OVA, αCD40 and αCD40-mSAs in cross-sections of the dLNs (**Fig. 3A**). Notably, mice treated with αCD40-mSAs-Bio-OVA exhibited stronger fluorescence signals of Bio-OVA in the dLNs, compared to those receiving Bio-OVA alone or a mixture of Bio-OVA and αCD40. The signal of αCD40-mSAs was also stronger than that of αCD40 and spatially overlapped with the Bio-OVA signal. Quantitative analysis revealed that, when normalized to tissue autofluorescence in naïve dLNs (set to 1 in the intensity ratio), the αCD40-mSAs-Bio-OVA vaccine group exhibited a 1.54-fold increase in Bio-OVA signal intensity. In contrast, the vaccine group receiving the Bio-OVA and αCD40 mixture showed no significant difference in Bio-OVA signal intensity compared to the Bio-OVA only vaccine group (**Fig. 3B**).

**Figure 3.**
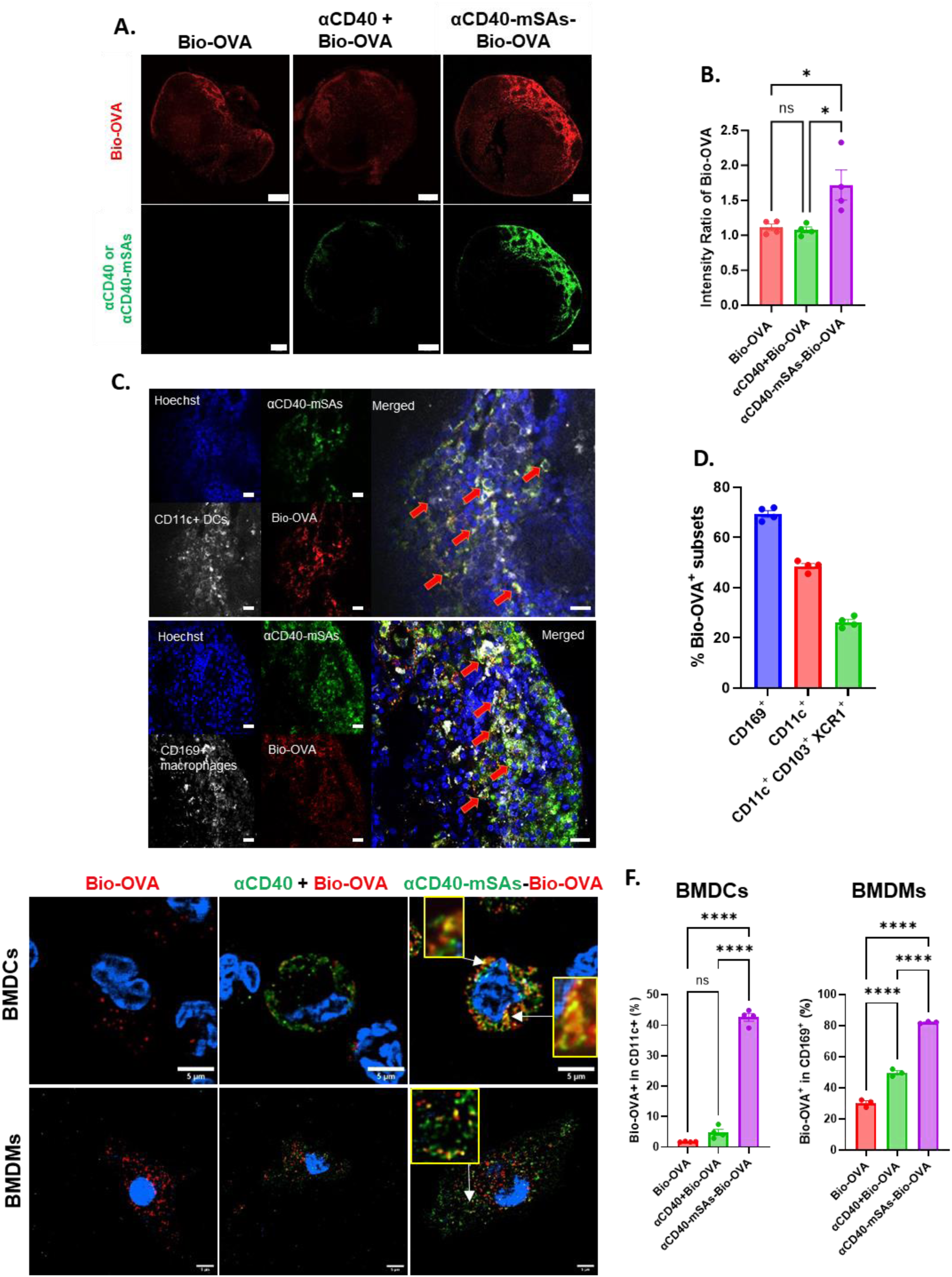
dLN-homing and APC-targeted intracellular delivery of neoantigen peptide by αCD40-mSAs. (A) Confocal microscope images showing the distribution of Bio-OVA-AF647 in mouse PO dLNs at 90 minutes after foot pad injection with peptide alone, αCD40-DL550, or αCD40-mSAs-DL550. Scale bar: 200 μm. (B) Quantification of Bio-OVA-AF647 signal intensity across different vaccination groups, normalized to the relative signal in PBS-injected controls. (C) High-resolution confocal images of PO dLNs confirming intracellular co-delivery of Bio-OVA-AF647 and αCD40-mSAs-DL550 in CD11c⁺ DCs (top) and CD169⁺ macrophages (bottom). Red arrows indicate individual cells showing overlapping signals of Bio-OVA and αCD40-mSAs. Scale bar: 20 μm. (D) Flow cytometry analysis identifying Bio-OVA⁺ APCs subsets, including DCs, macrophages, and cDC1s, in the PO dLNs after vaccination. (E) Super-resolution microscopy (Deep-SIM) of single BMDCs and BMDMs showing intracellular delivery of Bio-OVA-AF647 in different vaccine groups. Co-localization in single BMDCs and BMDMs, indicated by yellow-fluorescence in the inset boxes, reflects the overlap of Bio-OVA (red) and αCD40-mSAs (green) signals. Scale bar: 5 μm. (F) Flow cytometry results showing the frequency of CD11c⁺ BMDCs (left) and CD169⁺ BMDMs (right) that internalized Bio-OVA-AF647 in different vaccine groups. ns, not significant; \**p* < 0.05; \*\*\*\**p* < 0.0001.

To assess intracellular delivery of the neoantigen peptide into APCs, we performed high-resolution confocal microscopy of dLN sections stained with fluorescent anti-CD11c and anti-CD169 antibodies to identify DCs and subcapsular sinus (SCS) macrophages, respectively. Confocal microscopy revealed co-localization of Bio-OVA and αCD40-mSAs within a number of individual CD11c⁺ DCs and CD169⁺ SCS macrophages (**Fig. 3C**). Flow cytometry analysis of dLNs cells from mice receiving fluorescent αCD40-mSAs-Bio-OVA showed substantial uptake of OVA peptide by CD169⁺ SCS macrophages (∼70%) and CD11c⁺ DCs (∼50%) among the OVA⁺ lymphocyte population (**Fig. 3D**). Moreover, over half of the OVA⁺ CD11c⁺ DCs expressed CD103 and XCR1, markers of conventional type 1 DCs (cDC1s), which are specialized for antigen cross-presentation to CD8⁺ T cells (*45*). CD169⁺ SCS macrophages are also known to initiate cytotoxic T cell responses through interactions with DCs and other immune cells in the dLNs (*46*, *47*).

*In vitro* assays using bone marrow–derived dendritic cells (BMDCs) and macrophages (BMDMs) further confirmed intracellular co-delivery of Bio-OVA and αCD40-mSAs. Structured illumination microscopy (SIM) at single-cell resolution showed greater intracellular accumulation of Bio-OVA in αCD40-mSAs vaccine formulation-treated cells after incubation for 1 hour, compared to other groups (**Fig. 3E**). With higher spatial resolution, co-localization of Bio-OVA (red) and αCD40-mSAs (green) was evident within individual BMDCs and BMDMs, displayed as yellow signals in the zoomed-in insets (**Fig. 3E, right and Fig. S2**). Flow cytometry further confirmed significantly higher percentages of OVA⁺ CD11c⁺ BMDCs (>40%) and CD169⁺ BMDMs (>80%) following αCD40-mSAs-mediated delivery, relative to other groups (**Fig. 3F**). Enhanced internalization of αCD40-mSAs into both BMDCs and BMDMs was also observed (**Fig. S3**).

Together, these results demonstrate that neoantigen peptides loaded onto αCD40-mSAs *via* biotin-mSA interaction are efficiently and simultaneously delivered to dLNs proximal to the injection site and subsequently uptake by critical APC populations.

### Enhanced activation and antigen presentation of DCs by αCD40-mSAs

We examined the agonistic effects of αCD40-mSAs on the activation and antigen presentation of DCs *in vitro*. BMDCs were incubated with either αCD40 or αCD40-mSAs alone for 24 hours. PBS and LPS served as negative and positive controls, respectively. Compared to the PBS group, BMDCs treated with either αCD40 or αCD40-mSAs exhibited significantly higher expression of surface activation markers, including CD40 and CD80, as measured by flow cytometry **(Fig. 4A and 4B**). For CD40 detection, we used a neutralizing anti-CD40 antibody (clone 3/23) that does not interfere with or artificially enhance CD40 expression. This upregulation of CD40 expression in DCs was also confirmed in an *in vivo* study. Treatment with αCD40 alone increased both the absolute number and percentage of CD11c⁺CD40⁺ DCs in the dLNs (**Fig. S4**). This expansion of CD40⁺ DCs may contribute to a positive feedback loop, further enhancing immune responses upon repeated vaccinations. Activated BMDCs also secreted increased levels of the pro-inflammatory cytokine IL-12p40 **(Fig. 4C).** IL-12 production is a hallmark of DC maturation (*48*) and is known to promote the activation of CD8⁺ T cells (*49*). Interestingly, a direct comparison of BMDCs treated with αCD40 versus αCD40-mSAs showed no significant differences in surface activation marker expression or IL-12 production, suggesting that the monovalent streptavidin (mSA) component does not confer additional and noticeable immunostimulatory effects beyond the CD40 agonistic activity. Although streptavidin has been reported to exhibit immunostimulant properties in some vaccine strategies (*50*, *51*), it appears that mSA alone does not enhance DC activation in this context.

**Figure 4.**
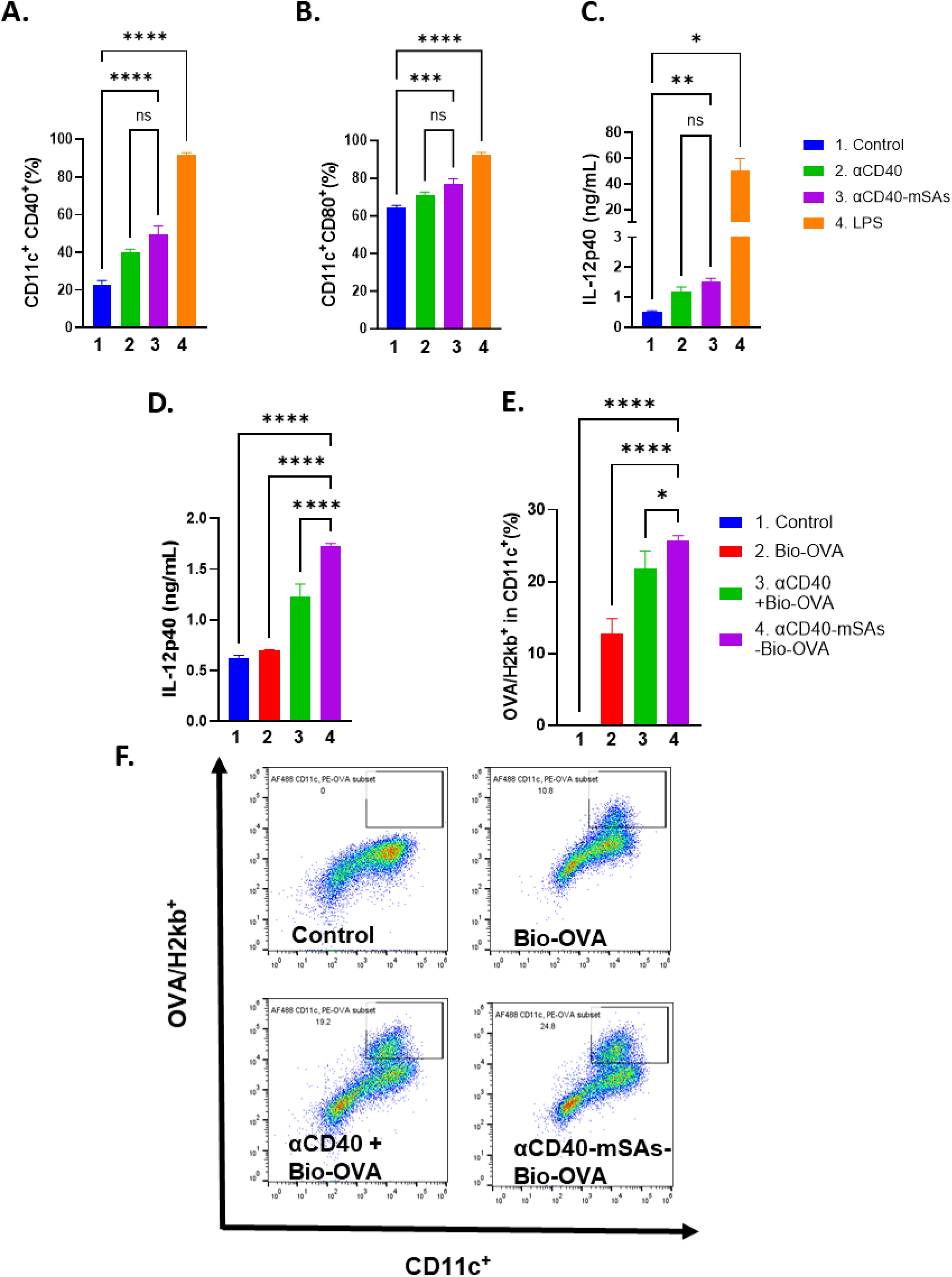
Agonistic effects of αCD40-mSAs on dendritic cell (DC) activation and antigen presentation. (A, B) Flow cytometry results showing expression of activation markers CD40 (A) and CD80 (B) on CD11c⁺ BMDCs after 24-hour incubation with control (no treatment), αCD40, αCD40-mSAs, or LPS. (C) ELISA results showing IL-12p40 levels in the culture supernatants of the treated BMDCs. (D) IL-12p40 levels in the culture supernatants of BMDCs treated with Bio-OVA vaccine formulations. (E) Percentage of CD11c⁺ BMDCs presenting the OVA epitope SIINFEKL (OVA/H2kb) after 24-hour incubation with different vaccine groups. (F) Representative flow cytometry plots of OVA epitope–presenting CD11c⁺ BMDCs treated with individual Bio-OVA vaccine groups. Data in (A-E) are presented as mean ± s.e.m. ns, not significant; *\*p* < 0.05; *\*\*p* < 0.01; *\*\*\*p* < 0.001; *\*\*\*\*p*<0.0001.

However, when BMDCs were incubated with αCD40-mSAs coupled with Bio-OVA, significantly higher IL-12 secretion was observed compared to the group treated with a mixture of αCD40 and Bio-OVA (**Fig. 4D**). Furthermore, antigen presentation was also significantly enhanced, reaching approximately 25%, compared to ∼20% in the αCD40 and Bio-OVA mixture group, and only ∼12% in the Bio-OVA alone group (**Fig. 4E**). This enhancement in antigen-presenting BMDC populations was clearly evident in the flow cytometry plots (**Fig. 4F**).

These results suggest that the effective intracellular delivery of neoantigen peptides by αCD40-mSAs primarily drives the superior DC activation and antigen presentation.

### Therapeutic vaccination using αCD40-mSAs against established primary tumors

We evaluated the therapeutic efficacy of the αCD40-mSAs vaccine formulation against established primary tumors using the syngeneic MC38 mouse colorectal tumor model. The neoantigenic peptide Adpgk (sequence provided in **Table S1**), which is endogenously expressed by MC38 cells, has been widely used as a model neoantigen in *in vivo* mouse tumor studies (*52*, *53*). MC38 tumor cells were inoculated subcutaneously into the flank, and tumor growth was monitored for 8 days post-inoculation. Once tumors reached a volume of over 100 mm³, tumor-bearing mice were vaccinated with different formulations, receiving a prime and two booster doses at one-week intervals (**Fig. 5A**). A PBS-injected group served as the control (Group 1, G1). Vaccine groups included Bio-Adpgk alone (G2), a mixture of αCD40 and Bio-Adpgk (αCD40 + Bio-Adpgk, G3), and Bio-Adpgk-loaded αCD40-mSAs (αCD40-mSAs-Bio-Adpgk, G4).

**Figure 5.**
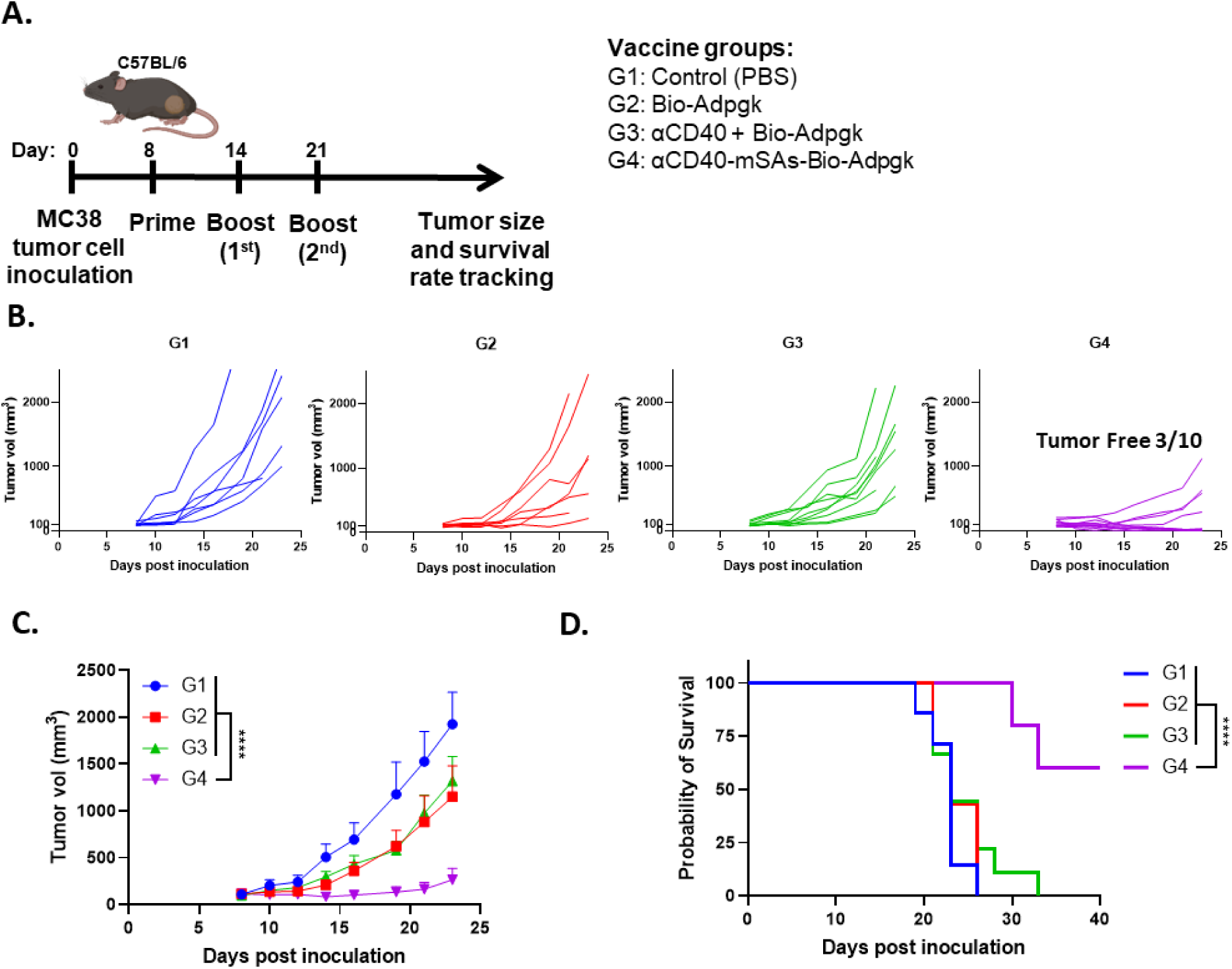
Therapeutic vaccine effects of neoantigen peptide-loaded αCD40-mSAs against established primary tumors. (A) Experimental timeline for therapeutic vaccination and monitoring of MC38 tumor growth and survival in tumor-bearing mice. Prime vaccination was performed on day 8 post-MC38 inoculation, when tumor volumes reached approximately 100 mm³. (B) Growth curves of individual MC38 tumors in mice treated with different vaccine groups (n = 7 for G1 and G2; n = 9 for G3; n = 10 for G4). In the G4 group (αCD40-mSAs-Bio-Adpgk vaccination), 3 out of 10 mice became tumor-free. (C) Average tumor growth curves across the different vaccine groups. Data are presented as mean ± s.e.m. (D) Kaplan–Meier survival curves for MC38 tumor-bearing mice following vaccination. *\*\*\*\*p* < 0.0001.

Tumor growth profiles from individual mice showed that vaccination with αCD40-mSAs-Bio-Adpgk (G4) significantly inhibited tumor growth compared to other groups (**Fig. 5B**). Although inter-mouse variability in tumor growth was observed within G4, 3 out of 10 mice achieved complete tumor regression during the 40-day monitoring period. In the processed tumor growth data (**Fig. 5C**), mean tumor volume curves and ANOVA analysis demonstrated significant tumor inhibition in G4, with tumor volumes markedly smaller than those of all other groups (*p* < 0.0001). While G2 (*p* = 0.0014) and G3 (*p* = 0.0293) also exhibited statistically significant tumor volume reductions compared to the G1 control group, the degree of inhibition was substantially lower than that observed in G4.

The rapid and aggressive growth of MC38 tumors caused mice in groups G1, G2, and G3 to reach humane endpoints within 23 days post-inoculation due to tumor ulceration and excessive tumor burden (**Fig. 5D**). In contrast, mice in G4 exhibited significantly prolonged survival. Kaplan-Meier survival analysis revealed a median survival of 23 days for mice in G1, G2, and G3. In contrast, the median survival of mice in the G4 group could not be determined, as a substantial number remained alive throughout the tumor monitoring period. Log-rank test results confirmed that mice in G4 showed significantly improved survival compared to those in all other groups (*p* < 0.0001), while no significant differences were observed among G1, G2, and G3. These results demonstrate the superior therapeutic efficacy of neoantigen peptide vaccination using αCD40-mSAs against established primary tumors.

### Prophylactic effect of αCD40-mSAs vaccination on tumor development

Next, we evaluated the prophylactic effect of vaccination on tumor development. Mice received a prime and two booster doses of the different vaccine formulations *via* footpad injection at 7-day intervals. Subsequently, MC38 tumor cells were inoculated into the flank (contralateral to the injection site) of the vaccinated mice (**Fig. 6A**). Mice vaccinated with αCD40-mSAs-Bio-Adpgk (G4) exhibited delayed tumor onset and significantly suppressed tumor growth up to day 24 post-inoculation compared to the other groups, as shown in individual tumor volume tracking (**Fig. 6B**). Representative images of mice from each group clearly showed that the smallest tumor was observed in the mouse from G4 compared to those from the other groups (**Fig. 6C**). The mean tumor growth curves further highlighted the delayed tumor progression in G4 relative to the PBS control group (G1) (*p* = 0.0136, **Fig. 6D**). Comparison between G4 and G1 did show statistically significant difference. Additionally, mice vaccinated with αCD40-mSAs-Bio-Adpgk (G4) demonstrated prolonged survival (**Fig. 6E**). The median survival times for G1, G2, G3, and G4 were 19, 26, 24, and 28 days, respectively. Log-rank test analysis confirmed that G4 mice had significantly improved survival compared to G1 (*p* = 0.0007) and G2 (*p* = 0.0313), whereas the difference between G4 and G3 was not statistically significant.

**Figure 6.**
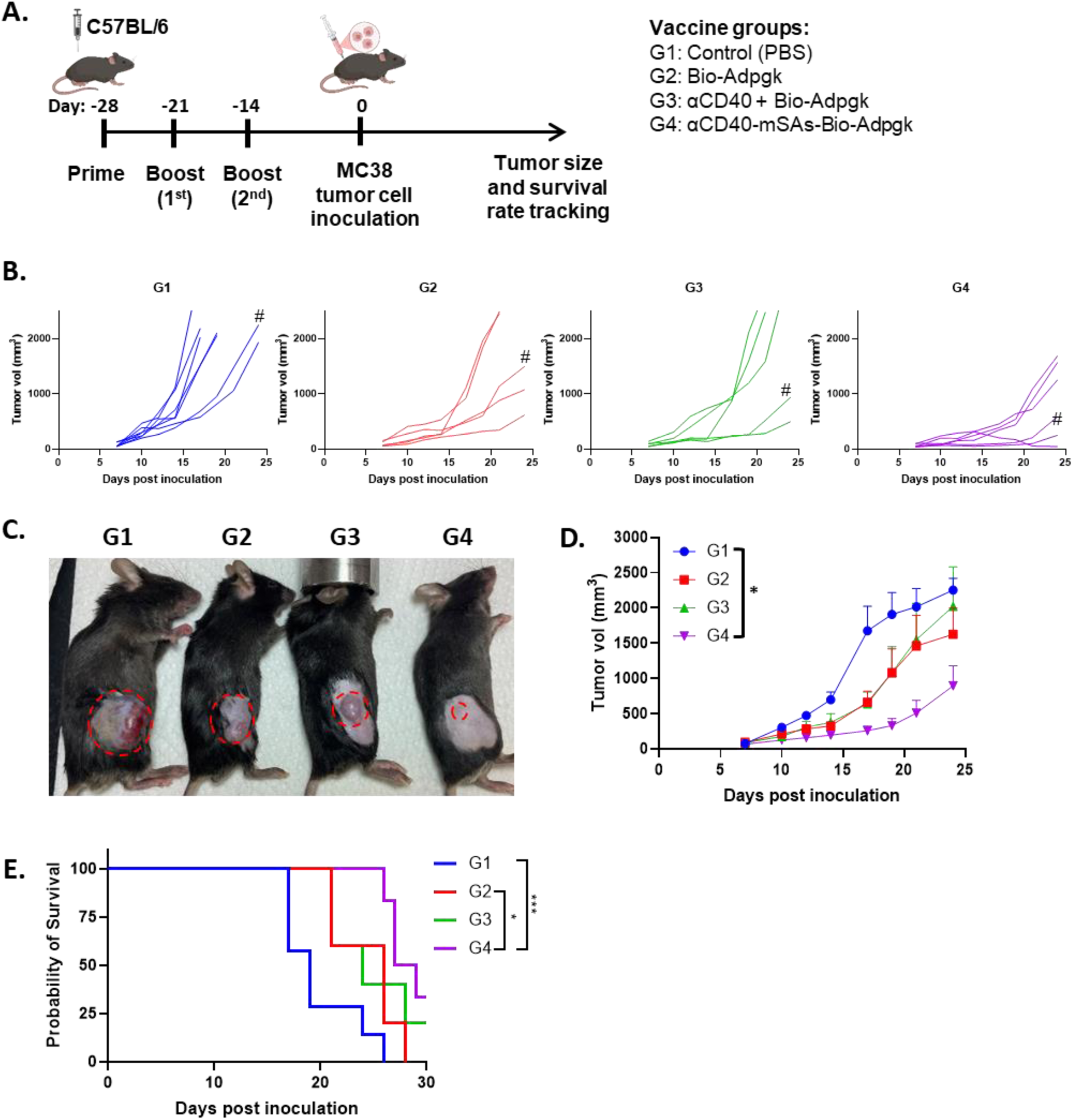
Prophylactic efficacy of neoantigen peptide-loaded αCD40-mSAs against tumor development. (A) Experimental timeline for prophylactic vaccination and monitoring of MC38 tumor development and survival in mice. (B) Growth curves of individual MC38 tumors in mice pre-treated with different vaccine groups (n = 7 for G1; n = 5 for G2; n = 5 for G3; n = 6 for G4). (C) Photograph of representative MC38 tumor-bearing mice from each vaccine group on day 21. The tumor growth curves of these individual mice are marked with ‘#’ in (B). (D) Average tumor growth curves across the different prophylactic vaccine groups. Data are presented as mean ± s.e.m. (E) Kaplan–Meier survival curves for pre-vaccinated mice following MC38 tumor inoculation. *\*p* < 0.05; *\*\*\*p* < 0.001.

### Induction of cancer neoantigen-specific effector cytotoxic T cell response by αCD40-mSA vaccination

To explore the immunological mechanisms underlying the prophylactic and therapeutic efficacy of αCD40-mSAs vaccination, we assessed the cytotoxic T cell response to the MC38 cancer neoantigen (Adpgk) following vaccination. Two weeks after a prime and two booster doses administered at 7-day intervals to naïve mice, dLNs and spleens were harvested for analysis of CD8⁺ T cell immunity (**Fig. 7A**). MC38 neoantigen-specific CD8⁺ T cells were identified in both dLNs and spleens using an H2-Db/MC38 (Adpgk, ASMTNMELM) MHC class I tetramer (**Fig. 7B**). A significantly higher percentage of MC38-specific CD8⁺ T cells was observed in both dLNs and spleens of mice vaccinated with αCD40-mSAs-Bio-Adpgk (G4) compared to the other vaccine groups (**Fig. 7C**). The induction of MC38 neoantigen-specific CD8⁺ T cells in both dLNs and spleens indicates the development of systemic adaptive anti-MC38 tumor immune responses. In addition, the frequency of CD8⁺ T cells producing TNF-α was elevated in both dLNs and spleens in the G4 group (**Fig. 7D**). TNF-α is a key immunostimulatory cytokine indicative of cytotoxic T cell activation. Notably, the increase in TNF-α-expressing CD8⁺ T cells was particularly pronounced in the spleen, with statistically significant differences observed between G4 and all other groups. In the dLNs, although the difference between G3 and G4 did not reach statistical significance, significant differences were observed between G4 and both G1 and G2. Collectively, these data indicate that αCD40-mSAs vaccination induces robust cancer neoantigen–specific adaptive T cell responses, leading to both therapeutic and prophylactic effects against tumor development and progression.

**Figure 7.**
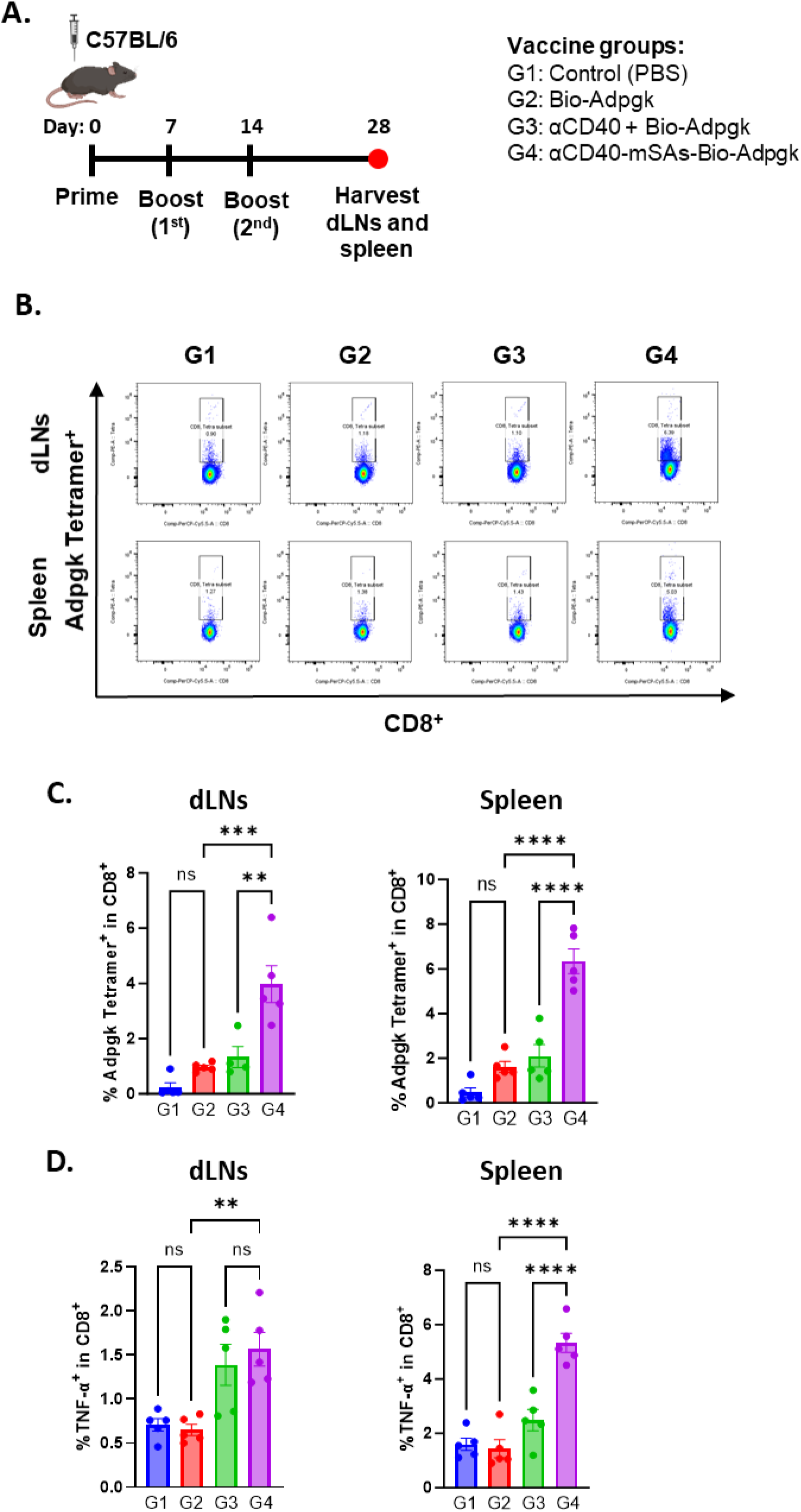
Enhanced induction of cancer neoantigen-specific CD8⁺ T cells and effector CD8⁺ T cell responses by vaccination with αCD40-mSAs. (A) Experimental timeline of vaccination and tissue collection. (B) Representative flow cytometry plots showing Adpgk tetramer^+^ CD8⁺ T cells in the dLNs and spleen. (C) Frequency of Adpgk tetramer^+^ CD8⁺ T cells in the dLNs and spleen. (D) Frequency of TNF-α secreting effector CD8⁺ T cells in the dLNs and spleen. ns, not significant; *\*\*p* < 0.01; *\*\*\*p* < 0.001; *\*\*\*\*p* < 0.0001.

## Discussion

We demonstrate that targeted delivery of neoantigen peptides is essential for effective cancer vaccine therapy. In particular, co-delivering the peptide with a molecular adjuvant directly into individual APCs within dLNs is crucial for inducing robust neoantigen-specific CD8⁺ T cell responses and suppressing tumor development and progression.

Most current clinical cancer vaccine trials use simple formulations consisting of mixtures of neoantigen peptides and adjuvants (*4*, *54*). Although comprehensive pharmacokinetic and biodistribution studies are lacking for these formulations, it is generally believed that the peptide and adjuvant are randomly and independently distributed after injection due to their distinct physicochemical properties (*55*). This uncontrolled delivery to APCs and dLNs can result in suboptimal vaccine efficacy and increase the risk of off-target effects and adverse immune responses (*16*, *56*). To address these limitations, it is essential to develop an optimized delivery system that ensures efficient and coordinated delivery of neoantigens and molecular adjuvants to APCs in dLNs.

Our engineered αCD40-mSAs construct is designed to address this key challenge in cancer vaccine delivery by leveraging two unique properties of an anti-CD40 agonistic IgG antibody: its natural dLN-homing capability and its molecular adjuvant activity. Endogenously produced IgG molecules circulate through the blood and lymphatic systems to capture antigens. Upon capturing antigens, antibody-antigen complexes drain into the dLNs, where they are taken up by APCs such as dendritic cells and macrophages, initiating innate and adaptive immune responses (*57*). Exogenously administered IgG antibodies demonstrate rapid drainage into nearby dLNs, primarily due to their large molecular weight (>84 kDa), which favors lymphatic over blood capillary absorption (*39*, *58*). Anti-CD40 agonistic antibodies have been widely used as molecular adjuvants that specifically target CD40-expressing APCs. Upon binding to CD40 on the plasma membrane of APCs, engagement of Fcγ receptors on neighboring immune cells facilitates CD40 crosslinking, which is essential for activating the CD40 signaling pathway (*34*, *59*, *60*). This crosslinking also promotes receptor-mediated endocytosis of the CD40–antibody complex into the APCs (*61*, *62*). These combined features, including dLN-homing, APC-targeting, intracellular uptake, and immune activation, make an anti-CD40 agonistic antibody an ideal platform for vaccine delivery.

To enable co-delivery of neoantigen peptides with this molecular adjuvant, we engineered the anti-mouse CD40 IgG2a antibody to express monovalent streptavidin (mSA) at the C-terminus of each light chain. Advances in protein engineering have enabled the generation of stable mSA constructs with high biotin-binding affinity (*42*, *63*). Although the adjuvant properties of mSA itself are not fully characterized, native streptavidin has been reported to enhance immune responses in certain vaccine settings, functioning as a molecular adjuvant (*50*).

On the other hand, biotinylation techniques are well-established and applicable to a wide range of biomolecules, including peptides, small molecules, and proteins (*64*, *65*, *40*). Due to its small molecular weight (∼224 Da), biotin conjugation has minimal impact on the physicochemical properties of peptides and is also biocompatible for clinical applications (*41*).

Building on these translational advantages and technical feasibility, we incorporated both mSA and biotinylation into the design of our engineered anti-CD40 agonistic antibody-based vaccine platform. The fusion of mSA at the C-terminus of each light chain preserves the antibody’s antigen-binding capacity through its variable regions, as well as its interactions with Fcγ receptors and FcRn *via* the Fc domain, both of which are critical for effector function, cellular targeting, and intracellular neoantigen delivery (*66–68*). This αCD40-mSAs construct offers flexible loading capacity for any biotinylated neoantigen peptide identified from individual cancer patients, enabling rapid customization for personalized cancer vaccine therapy. Therefore, αCD40-mSAs serves as an optimal “all-in-one” vaccine delivery platform that integrates targeted co-delivery of neoantigen peptide-molecular adjuvant complexes to dLNs and APCs, while maintaining adaptability for personalized cancer vaccination.

After confirming the biophysical properties and binding affinities of the engineered αCD40-mSAs to both mouse CD40 and biotin, we employed advanced imaging methods to evaluate the effectiveness of our delivery approach (*69*, *70*). In biodistribution studies, we observed enhanced dLN-homing and APC-targeted delivery of neoantigen peptides using αCD40-mSAs. Notably, among the APC populations that internalized neoantigen peptides, we identified a substantial fraction of CD11c⁺CD103⁺XCR1⁺ cDC1 and CD169⁺ SCS macrophages, both of which are well-characterized as key players in initiating neoantigen-specific cytotoxic T cell responses (*45*, *46*). Furthermore, single-cell resolution microscopy in *in vitro* assays confirmed the intracellular co-delivery of both neoantigen peptides and αCD40-mSAs into the same individual APCs. These results support the αCD40-mSAs as an effective platform for targeted neoantigen peptide delivery.

We also confirmed the superior agonistic function of αCD40-mSAs in promoting DC activation and antigen presentation through *in vitro* studies. Importantly, CD40 pathway-targeted DC activation led to the upregulation of CD40 expression, potentially establishing a self-amplifying loop that could further enhance immune responses upon repeated αCD40-mSAs vaccination. *In vivo* vaccination with cancer neoantigen-loaded αCD40-mSAs demonstrated significant anti-tumor therapeutic and preventive effects, resulting in prolonged survival. Notably, complete tumor remission was observed in some cases of established tumors. T cell profiling of mouse dLNs and spleen following αCD40-mSAs vaccination revealed the generation of cancer neoantigen-specific effector cytotoxic T cell populations, indicating a systemic, vaccine-induced adaptive immune response against target tumors. These results highlight the importance of targeted co-delivery of neoantigens and molecular adjuvants for inducing potent cancer neoantigen-specific cytotoxic T cell responses.

Given the advantages of anti-CD40 agonistic antibodies in vaccination, various antigen-loading strategies have been explored, including chemical conjugation, non-covalent assembly, and genetic fusion with antigen sequences (*71–74*). Although these methods have shown promising vaccination outcomes in preclinical and clinical studies, these antibody-based vaccines often require lengthy synthesis processes and offer limited adaptability to diverse antigens, making them less suitable for custom formulations used in personalized cancer vaccination. A recent study engineered an anti-CD40 agonist antibody to express additional Fab arms at the C-terminus of the light or heavy chains (*75*). These Fab arms in the engineered CD40 antibody were designed to bind a 9–amino acid (AA) peptide (p)-tag peptide. As a result, neoantigen peptides containing the p-tag sequence can be attached to the Fab arms, providing flexibility to load different neoantigen peptides onto the bispecific CD40 antibody system. Treatment with the bispecific CD40 antibody in combination with p-tagged neoantigen peptides induced enhanced neoantigen-specific immune responses compared to the group receiving non-p-tagged peptides. Despite demonstrating improved vaccination efficacy, the delivery mechanism of this bispecific CD40 antibody system, actually designed for co-delivery, remains to be elucidated. Although the shortened 9-AA p-tag derived from the original 18-AA sequence (*76*) exhibits sufficient binding affinity to the anti-p-tag_9AA_ Fab, it remains unclear whether this interaction is stable *in vivo* to support reliable co-delivery of both the neoantigen and CD40 antibody to individual APCs in the dLNs. Moreover, even with the shorter 9-AA p-tag, the required peptide extension may alter the biophysical properties of certain short neoantigen peptides, potentially affecting their pharmacokinetics and immunogenicity. In contrast, our mSA-biotin system provides a simpler yet sufficiently stable interaction for co-delivery into dLNs and APCs, as demonstrated in this study. Furthermore, the small molecular weight of biotin and the versatility of its conjugation chemistry enable biotinylation of validated SLPs or short epitope peptides without significantly altering their biophysical properties, while allowing efficient loading onto the αCD40-mSAs platform.

Despite enhanced neoantigen delivery, the anti-tumor therapeutic and preventive efficacy of αCD40-mSAs vaccine did not result in complete tumor regression in all treated mice. As a potential future direction, combining αCD40-mSAs cancer vaccination with immune checkpoint blockade therapy may improve *in vivo* outcomes, as checkpoint blockade can prevent T cell exhaustion and further enhance vaccine-induced immunity (*77*, *78*). Additionally, a heterologous vaccination strategy using a cocktail of αCD40-mSAs loaded with different neoantigen peptides could help prevent immune escape due to antigen loss, which is a strategy commonly employed in peptide-based cancer vaccine clinical trials (*10*, *11*).

In conclusion, this study demonstrates the immunopharmacological advantages of αCD40-mSAs as an optimal neoantigen peptide delivery platform for cancer vaccination, including dLN-homing, APC-specific intracellular delivery, and immunostimulation. The simple and stable ‘plug & play’ peptide-loading feature of αCD40-mSA holds strong potential to enhance the effectiveness of personalized cancer vaccination and improve clinical outcomes through precise and efficient neoantigen delivery.

## Materials and Methods

### Mice

C57BL/6 male mice (6-8 weeks old) were purchased from Inotiv (formerly Envigo) and housed under standardized conditions in the animal facilities at the University of Illinois Chicago. All animal experiments were approved by and conducted in accordance with the ethical guidelines of the Institutional Animal Care and Use Committee (IACUC) of the University of Illinois Chicago.

### Cell line

MC38 (colorectal cancer) cells were purchased from ATCC (VA, USA) and maintained *via* the ATCC-recommended culture methods. Cells were cultured in Dulbecco’s Modified Eagle Medium (DMEM; Gibco, USA) supplemented with 10% fetal bovine serum (FBS; Corning, USA), 2 mM L-glutamine (Gibco, USA), 1× non-essential amino acids (NEAA; Ginco, USA), 1 mM sodium pyruvate, and 1% penicillin–streptomycin at 37 °C in a humidified incubator with 5% CO₂.

### Peptides

Biotinylated synthetic long peptides (SLPs) containing the sequence of OVA (SIINFEKL) or Adpgk (ASMTNMELM) (*52*), with or without fluorophore conjugation, were synthesized by LifeTein LLC (**Table S1**). All peptides had high purity (>95%), and their sequences were confirmed by high-performance liquid chromatography and mass spectrometry (HPLC-MS). Peptides were initially reconstituted at 1 mM in sterile water and subsequently diluted to working concentrations in phosphate-buffered saline (PBS).

### Synthesis of recombinant αCD40 and αCD40-mSAs

Codon-optimized sequences for the variable light (VL) and variable heavy (VH) chains of the anti-mouse CD40 agonistic antibody (clone FGK4.5) as well as monovalent streptavidin (mSA) (*63*) were synthesized (ThermoFisher Scientific, Waltham, MA) and cloned into the pTRIOZ-mIgG2a/κ vector (InvivoGen, CA). Plasmids encoding the αCD40 and αCD40-mSAs antibodies, pTRIOZ-mIgG2a/mk-mαCD40 and pTRIOZ-mIgG2a/mk-mαCD40-mSAs, respectively, were transfected into ExpiCHO-S cells using the ExpiCHO transfection kit (ThermoFisher Scientific, Waltham, MA) according to the manufacturer’s instructions. ExpiCHO-S cells were cultured in ExpiCHO Expression Medium (ThermoFisher Scientific) on a shaker set to 120 rpm, in an incubator at 32 °C with 8.0% CO_2_. Cells were collected 12 days post-transfection by centrifugation at 4,000 × g and 4 °C for 20 minutes. The antibody supernatant was passed through a 0.22-µm filter and neutralized with 10× PBS buffer (Cytiva, Marlborough, MA). The antibodies were purified using the AKTA Start protein purification system and HiTrap MabSelect SuRe protein A column (Cytiva). The purified antibodies were concentrated and buffer exchanged with PBS (Cytiva) using Vivaspin® Turbo (MWCO 10k Da; Sartorius, Göttingen, Germany). The antibody concentration was determined by UV absorbance at 280 nm. The purity of the synthesized antibodies was determined by SDS-PAGE analysis using gels and Coomassie blue staining (Bio-Rad), performed according to the manufacturer’s protocol. Gel band images were acquired using Odyssey CLx infrared imaging system (LI-COR Biosciences, Lincoln, NE). Endotoxin levels in the antibody solutions were measured using the Pierce^TM^ Chromogenic Endotoxin Quant Kit (ThermoFisher Scientific). The antibody solutions contained less than 0.5 EU endotoxin per mg of antibody.

### Enzyme-linked immunosorbent assay (ELISA)

ELISAs were performed to assess the binding of αCD40 and αCD40-mSAs to mouse CD40 protein and biotin. For the ELISA assay measuring binding to mouse CD40, Pierce Ni-NTA– coated 96-well plates (Thermo Fisher Scientific) were coated with mouse CD40-His protein (1 μg/mL, AcroBiosystems, Newark, DE) for 1 hour at RT, followed by incubation with αCD40 or αCD40-mSAs antibodies (0.0125 to 4 μg /mL) for 1 hour RT. After washing with PBS, HRP-conjugated anti-mouse IgG secondary antibodies (dilution of 1:10000, Thermo Fisher Scientific) were incubated for 30 minutes. Upon addition of TMB substrate (ThermoFisher Scientific), the bound αCD40 or αCD40-mSAs antibody was quantified by measuring OD_450_ value using a Synergy LX multi-mode reader (Agilent, Santa Clara, CA).

For the mouse CD40/CD40L blockade assay, Pierce Ni-NTA–coated 96-well plates (Thermo Fisher Scientific) were first coated with mouse CD40-His protein (0.1 μg/well), followed by incubation with mouse CD40L-hFc protein (human Fc–conjugated; 1 μg /mL, AcroBiosystems). After washing with PBS, αCD40 or αCD40-mSAs antibody (0.0125 to 4 μg /mL) was added to assess blocking activity. Bound CD40L-hFc was then detected using HRP-conjugated anti-human IgG Fc–specific secondary antibody (80 ng/mL, Thermo Fisher Scientific). Upon addition of TMB substrate, signal intensity was measured at OD₄₅₀ using a Synergy LX multi-mode reader (Agilent, Santa Clara, CA).

For biotin binding, αCD40 or αCD40-mSAs (0.2 μg/well) was first captured in anti-mouse IgG antibody-coated 96-well plates (ThermoFisher Scientific) and incubated for 1 hour at RT. After washing with PBS, serial dilutions of biotin-HRP (0.064-8 ng/mL; ThermoFisher Scientific) were added and incubated for 1 hour at RT. TMB substrate (ThermoFisher Scientific) was added for detection, and OD_450_ was measured.

### SDS-PAGE

The molecular weight and expression of the mSAs of synthesized αCD40-mSAs were confirmed using SDS-PAGE analysis in reducing and non-reducing conditions. A total of 1 μg of the purified αCD40 or αCD40-mSAs was mixed with 4× LDS sample buffer (ThermoFisher Scientific) for both reducing and non-reducing conditions. PNGase F (New England Biolabs, MA) was used to remove glycosylation of the antibody. The samples were run on a MiniPROTEAN® TGXTM Precaset Gel 4-15% (Bio-Rad) and Coomassie blue staining were performed according to the manufacturer’s protocol (Bio-Rad). Gel band images were acquired using Odyssey CLx infrared imaging system (LI-COR Biosciences, Lincoln, NE)

### Mass photometry (MP) analysis

Mass photometry (MP) measurements were performed using a Refeyn Two MP instrument. First, a drop of immersion oil (Thorlabs, cat. MOIL-30) was applied to the objective. A clean glass slide (GLW Storing Systems, cat. ZK25) was then positioned atop the oil along with a reusable culture well gasket (3 mm diameter × 1 mm depth; Sigma, cat. GBL103250-10EA). Next, 20 µL of PBS was added into the gasket, and the objective was carefully adjusted and focused on the solution. The instrument was calibrated using a protein standard in PBS, consisting of Thyroglobulin (Tg: 330 and 660 kDa as monomer and dimer), and β-amylase (BAM: 56, 112, and 224 kDa as monomer, dimer, and trimer) in PBS. For each measurement, 10 µL of buffer was first added to the well for focusing, after which 10 µL of the sample was introduced and gently mixed. Data acquisition started immediately after the mixture, with movies recorded for 120 seconds at 100 frames per second under standard settings. The resulting mass photometry data were analyzed using DiscoverMP (Refeyn Ltd.).

### Fluorophore polarization (FP) analysis

Interactions with biotin were assessed by titrating αCD40-mSAs and streptavidin (Millipore-Sigma, cat. 189730) and over a concentration range of 0.007 to 2000 nM in PBS using fluorophore polarization (FP) analysis. Biotin-FITC (Millipore-Sigma, cat.53608) was prepared at a final concentration of 20 nM in PBS after initial reconstitution in DMSO. All assays were performed in 384-well black flat-bottom plates (Corning) with a final volume of 40 µL per well. FP measurements were acquired using a Tecan Infinite F200 Pro at the UIC High Throughput Screening Core, where the perpendicular emission signal was subtracted from the parallel emission signal to yield the millipolarization (mP) values. Excitation and emission wavelengths were set to 485 nm and 535 nm, respectively. K_d_ values were determined by curve fitting the dose-dependent FP signal as a function of protein concentration using Prism 10 (GraphPad Software).

### Surface plasmon resonance (SPR) analysis

SPR experiments were performed on a Biacore T200 instrument (GE Healthcare, Sweden). CM5 sensor chips, along with the amine coupling kit and regeneration solutions, were obtained from Cytiva. Mouse CD40/TNFRSF5 protein (Sino Biological, cat. 50324-M08H) was immobilized onto the CM5 sensor chip *via* standard amine coupling chemistry until a level of approximately 450 response units (RU) was reached. Following immobilization, three analytes were evaluated: a commercial anti-CD40 antibody that shares the same CDR sequence with Rat IgG construction (Clone FGK4.5, Bioxcell), synthesized αCD40, and αCD40-mSAs. Each analyte was injected in a series of five concentrations (ranging from 0.37 to 30 nM) prepared by threefold serial dilutions. During each cycle, the analyte was allowed to associate with the immobilized ligand for 90 seconds, followed by a 360-second dissociation phase. All injections were conducted at a flow rate of 30 µL/min and maintained at 25 °C. Between cycles, the sensor surface was regenerated by injecting 10 mM glycine-HCl (pH 2 or 3, as optimized for each analyte) at a flow rate of 30 µL/min. Sensorgrams were recorded and subsequently analyzed using Biacore Insight Evaluation Software (version 3.0.12), with kinetic parameters determined by fitting the data to a 1:1 Langmuir binding model.

### *In vivo* assessment of antigen-specific cytotoxic T cell response

C57BL/6 mice were randomly assigned to four vaccine groups (n = 5 per group): Group 1 (Control): received PBS only. Group 2 (Bio-OVA): received 3.32 μg of biotinylated ovalbumin (OVA) peptide (40 μM). Group 3 (αCD40+Bio-OVA): received 3.32 μg of biotin-OVA peptide (40 μM) combined with 90 μg of αCD40 (20 μM). Group 4 (αCD40-mSAs-Bio-OVA): received 3.32 μg of biotin-OVA peptide (40 μM) loaded onto 108 μg of engineered αCD40-mSAs (20 μM). Before injection, αCD40 and αCD40-mSAs were incubated with the Bio-OVA peptide for 30 minutes at RT. All treatments were administered *via* a single footpad injection in the right side of hindlimb of each mouse in a volume of 30 μL. Spleens and lymph nodes, including popliteal (PO), inguinal (IN), and axillary (AX) lymph nodes from both sides (total of 6 per mouse), were harvested seven days post-vaccination. Single-cell suspensions were prepared by mechanical dissociation using a mesh filter with a pore size of 40 μm (Greiner, cat.542140). The cells underwent red blood cell lysis using ice-cold RBC Lysis Buffer (Roche, cat. 11814389001) for 5 minutes, followed by washing in cold PBS. Then, the cells were stained with fluorescent anti-CD3, CD8, and CD44 antibodies (antibody details in **Table S2**) and OVA epitope-specific PE-labeled MHC I tetramer (obtained from the NIH Tetramer Core Facility at Emory University). For flow cytometry analysis, the gating strategy is detailed in **Fig. S5A**. The frequency and absolute number of OVA-specific cytotoxic T cells (OVA tetramer⁺ CD44⁺ CD8⁺ CD3⁺) were quantified in both dLNs and spleens. Detailed flow cytometry procedures and analyses are described in the ‘Flow cytometry analysis’ section below.

### *In vivo* biodistribution in dLNs

#### Confocal Microscopy

αCD40 and αCD40-mSAs were conjugated with DyLight550 (DL550) NHS Ester (Thermo Fisher Scientific) by overnight incubation at a 1:20 molar ratio (antibody:dye). Unconjugated dye was removed using a Zeba Spin Desalting Column (Thermo Fisher Scientific). The fluorescently labeled antibodies were protected from light and stored at 4 °C until use. Mice (n=3 per group) were injected in the right hindlimb footpad with a 30 μL vaccine solution containing either biotin-OVA conjugated with Alexa Fluor 647 (Bio-OVA-AF647) alone (10 μM), a mixture of Bio-OVA-AF647 (10 μM) and αCD40-DL550 (5 μM), or Bio-OVA-AF647 (10 μM) loaded onto αCD40-mSAs-DL550 (5 μM). At 90 minutes post-injection, popliteal (PO) dLNs on the injected side were harvested, embedded in 2% (w/v) agarose gel, and sectioned into 200 μm slices using a vibratome (VT1200S, Leica Biosystems). After fixation in 2 % para-formaldehyde (PFA) for 10 minutes at RT and washing in PBS three times, the dLNs slices were stained with Hoechst 34580 (at 0.1 mg/mL) and either anti-CD11c-AF488 (at 0.2 mg/mL) or anti-CD169-AF488 (at 0.02 mg/mL). Details of the antibodies used for microscopy are described at **Table S3**. Whole cross-sectional and high-resolution images of the dLNs were acquired using a Zeiss Laser Scanning Confocal Microscope (LSM 710) with a 20× and 63× objectives and 488/556/633 nm excitation lasers and corresponding emission filter settings. Quantitative fluorescence intensity analysis was performed using ImageJ or Fiji software by measuring the signal intensity in each image and normalizing it to the fluorescence intensity observed in PBS-injected control dLNs.

#### Flow cytometry

Mice were injected in the footpad with Bio-OVA-AF647 (20 μM) loaded onto αCD40-mSAs (10 μM). At 90 minutes post-injection, popliteal (PO) dLNs on the injected side were harvested and mechanically dissociated into single cells. Cells were initially incubated with an Fc block antibody to prevent non-specific binding, and stained with anti-CD11c-BV421, CD169-AF488, and CD103-AF700, and XCR1-PE/Dazzle594 in 100 µL cell staining buffer. The percentage of CD11c⁺ cells, CD169⁺ cells, and CD11c⁺ CD103⁺ XCR1 ⁺ cells presenting the OVA peptide was subsequently quantified by flow cytometry. Gating strategy for flow cytometry analysis is in **Fig. S5B**. Detailed flow cytometry procedures and analyses are described in the ‘Flow cytometry analysis’ section below and the antibody product and dilution details in **Table S2**.

### Intracellular vaccine delivery in APCs

#### Flow cytometry

Bone marrow-derived dendritic cells (BMDCs) were generated from femurs and tibias of C57BL/6 male mice (6-8 weeks old). Following euthanasia, bone marrow was flushed from the bones and cultured in RPMI-1640 medium supplemented with 10% fetal bovine serum (FBS) and 20 ng/mL recombinant mouse GM-CSF (R&D Systems, cat. 415-ML) at 37 °C in a humidified atmosphere containing 5% CO₂. On days 6-7, loosely adherent and non-adherent cells exhibiting dendritic morphology were harvested and used for subsequent *in vitro* experiments. Bone marrow-derived macrophages (BMDMs) were generated from isolated bone marrow of C57BL/6 male mice (6-8 weeks old) and cultured in macrophage culture medium consists of DMEM/F12 medium supplemented with 10% fetal bovine serum (FBS), 20 ng/mL recombinant mouse macrophage colony-stimulating factor (M-CSF) (R&D Systems), and 2 mM glutamax supplementary. BMDCs and BMDMs were incubated with PBS, Bio-OVA-AF647 (1 μM), αCD40 (500 nM) + Bio-OVA-AF647 (1 μM), αCD40-mSAs (500 nM)-Bio-OVA-AF647 (1 μM) at 37 ℃ for 1 hour, and stained with anti-CD11c-AF488 for 30 minutes for BMDCs and anti-CD169-AF488 for BMDMs. The percentage (%) of CD11c⁺ or CD169⁺ cells presenting OVA peptide and antibody was subsequently quantified by flow cytometry (CytoFlex). Detailed flow cytometry procedures and analyses are described in the ‘Flow cytometry analysis’ section below and the antibody product and dilution details in **Table S2**.

#### DeepSIM super-resolution microscopy

BMDCs and BMDMs treated with the vaccine formulations were prepared using the same methods described above, followed by fixation and staining with Hoechst 34580. The DeepSIM images were acquired using a CrestOptics DeepSIM Lattice SIM super-resolution module equipped on a Nikon Ti2-E inverted microscope. Imaging was performed using a Nikon 60x/1.42 oil immersion objective and a Kinetix sCMOS camera (Serial Number: A23H723002). The 405 nm, 545 nm, and 637 nm laser lines from a Lumencor CELESTA Light Engine were used with a power percentage of 5%, 70%, and 70%, out of 621 mW, 708 mW, and 779 mW at 100% intensity, respectively. The power at 100% is from the CELESTA Light Engine output. The camera exposure times were set to 10 ms for the 405 nm, 300 ms for the 545 nm, and 300 ms for the 637 nm laser excitation. Images were acquired using the Deep imaging mode. All imaging was performed at room temperature. Raw images were reconstructed using Nikon Elements Advanced Research (Version: 5.42.04) software with the DeepSIM module based on the Richardson-Lucy Deconvolution algorithm, resulting in a pixel size of 54 nm in the reconstructed image.

### *In vitro* evaluation of agonistic effects in DCs

One million BMDCs were incubated with either αCD40 (30 nM), αCD40-mSAs (30 nM), or LPS (10 ng/mL) for 24 hours at 37 ℃ in a 24 well plate (Corning, USA). The BMDCs were stained with fluorescent anti-CD11c, CD80, and CD40 (neutral, clone 3/23) antibodies for flow cytometry analysis of DC activation and maturation. Immunocytokine, IL-12p40, were measured in culture media supernatant using an ELISA kit (Biolegend, cat. 431604) by following manufacturer instructions. For antigen presentation assays, BMDCs were incubated with either Bio-OVA alone (10 μM), a mixture of Bio-OVA (10 μM) and αCD40 (5 μM), or Bio-OVA (10 μM)-loaded αCD40-mSAs (5 μM) for 24 hours at 37 ℃. Antigen presentation was detected by staining with an anti-H-2Kᵇ-SIINFEKL antibody (BioXcell), which specifically recognizes the SIINFEKL epitope (OVA₂₅₇-₂₆₄) bound to MHC class I (H-2Kᵇ). The percentage of CD11c⁺ BMDCs presenting the OVA epitope was subsequently quantified by flow cytometry. The BMDC gating strategy for flow cytometry analysis is in **Fig. S6** and detailed flow cytometry procedures and analyses are described in the ‘Flow cytometry analysis’ section below and the antibody details in **Table S2**.

### *In* vivo therapeutic vaccine effect on established tumors

C57BL/6 male mice (6-8 weeks old) were subcutaneously inoculated in the shaved right hind flank with 2.5 × 10⁵ MC38 cells suspended in 100 µL of a 1:1 mixture of culture media and Matrigel (Corning, cat. 354230). Once a tumor volume reached to 100 mm³ at 8 days post-inoculation, mice were randomized into four groups for vaccination. Vaccine groups included: (Group 1) control group injected with PBS, (Group 2) Bio-Adpgk peptide alone (40 μM), (Group 3) a mixture of Bio-Adpgk (40 μM) and αCD40 (20 μM), and (Group 4) Bio-Adpgk (40 μM) loaded on αCD40-mSAs (20 μM). The prime and two boosting vaccinations (each dose in 30 μL) in a week interval performed by food pad injection. For the vaccination with Group 3 and 4, Bio-Adpgk was pre-incubated with αCD40 or αCD40-mSAs for 30 minutes before injection. Tumor dimensions (length and width) were measured every 2 or 3 days using an electronic caliper, and tumor volume was calculated using the formula: Volume = (length × width²)/2. Mice were euthanized upon reaching humane endpoints, defined as a tumor volume exceeding 2000 mm³ or ulceration affecting more than 30% of the tumor area. Tumor growth was tracked until day 26, and survival was monitored until day 40.

### *In vivo* prophylactic vaccine effects on tumor development

C57BL/6 male mice (6-8 weeks old) were vaccinated with the same groups and regimen described above, using one prime and two booster doses administered at one week intervals by right hind foot pad injection. At two weeks following the final vaccination, 2.5 × 10⁵ MC38 cells were inoculated into the left hind flank of the vaccinated mice subcutaneously. Tumor volume was measured by the same way described above for 24 days post inoculation. The survival period was tracked until 30 days post inoculation.

### Induction of cancer antigen-specific effector cytotoxic T cells

C57BL/6 male mice (6-8 weeks old) were vaccinated with the same groups and regimen described above in one prime and two boosting at a week interval. At 2-weeks post the final vaccination, the mice were sacrificed, and the spleens and dLNs (popliteal (PO), inguinal (IN), and axillary (AX) lymph nodes from both sides, total of 6 per mouse), were harvested. The spleens and dLNs were mechanically dissociated into single-cell suspensions, filtered through 40 µm cell strainers, and washed with ice-cold PBS. Cells underwent red blood cell lysis using ice-cold RBC Lysis Buffer (Roche, cat. 11814389001) for 5 minutes, followed by washing in cold PBS. All samples were re-suspended in flow cytometry staining buffer (BioLegend, cat. 420201) and stained with fluorescent anti-CD3, CD4, CD8, CD62L, CD44 antibodies for flow cytometry analysis. MC38 cancer-specific CD8⁺ T cells were identified using a PE-labeled MHC I tetramer specific to the Adpgk epitope (obtained from the NIH Tetramer Core Facility at Emory University). Intracellular cytokine expression of TNF-α in CD8^+^ cytotoxic T cells was measured by flow cytometry after staining with fluorescent anti-TNF-α antibody after cell membrane permeabilization using Fixation/Permeabilization Kit (BioLegend, cat. 426803). T cell profiling gating strategy for flow cytometry analysis is in **Fig. S7,** and detailed flow cytometry procedures and analyses are described in the ‘Flow cytometry analysis’ section below.

### Flow cytometry analysis

BMDCs, BMDMs, and single-cell suspensions from lymph nodes and spleens were prepared as described above. For surface staining, tetramer staining, and intracellular cytokine staining, cells were incubated with the appropriate fluorescence-conjugated antibodies. Detailed antibody dilution factors, product information, fluorochrome conjugates, and clone names are provided in **Table S2**. Cells (one million cells per sample) were resuspended in 100 µL PBS, and all subsequent staining procedures were performed in the dark. Initially, cells were stained with Live/Dead Zombie Violet Dye (BioLegend, cat. 423113) to assess viability, followed by washing and resuspension in 100 µL of flow cytometry staining buffer. Next, cells were incubated with TruStain Mouse FC Block (BioLegend, cat. 101320) for 10 minutes at room temperature to reduce nonspecific binding. For experiments requiring antigen-specific tetramer staining, tetramers (obtained from the NIH Tetramer Core Facility) were added immediately after Fc block incubation and incubated for 20 minutes before adding surface antibodies. Subsequently, the surface antibody cocktail was added and incubated for 30 minutes. For intracellular cytokine staining, cells underwent additional fixation and permeabilization using the Fixation/Permeabilization Kit (BioLegend, cat. 426803) following the manufacturer’s instructions. Intracellular staining for cytokines was performed by incubating the permeabilized cells with the relevant antibodies (product name and details provided in **Table S2)** for 30 minutes. After staining, cells were washed, resuspended in PBS, and stored at 4 °C until flow cytometry analysis. Single-stain compensation was carried out using Compensation Beads (BioLegend, cat.424601). Samples were acquired on different cytometers based on the specific assay: Gallios Flow Cytometer (Beckman Coulter, USA) for BMDC activation and antigen presentation assays, and *In vivo* biodistribution in dLNs studies, Cytoflex Flow Cytometer (Beckman Coulter, USA) for Intracellular uptake in BMDCs and BMDMs, and Cytek Aurora (Cytek Biosciences, USA) for T cell evaluation and cytokine profiling. For data acquired by Gallios and Cytofelx were compensated, gating, and analysis were conducted using FlowJo v10. For data acquired on the Cytek Aurora, spectral unmixing was performed using SpectroFlo software, and compensation and further analysis were conducted using FlowJo v10.

### Statistical analysis

Group mean comparisons were performed using GraphPad Prism software, V.10.4.1. One and two-way Analysis of variance (ANOVA) with Tukey’s multiple-comparison test was used for a comparison of multiple group’s means. Kaplan-Meier curves for survival analysis were conducted using a Mantel-Cox log-rank test. For all comparisons, *p* values ≤0.05 were considered statistically significant. Significance levels are indicated in the figures as follows: *p* > 0.05, no significance, n.s.; \**p* < 0.05; \*\**p* < 0.01; \*\*\**p* < 0.001; \*\*\*\**p* < 0.0001

## Acknowledgments

The SPR, ITC, and MP analysis were performed using the instruments in the Biophysics Core in Research Resources Center of University of Illinois Chicago. We acknowledge UIC High Throughput Screening Core director Dr. Kiira Ratia for assistance and guidance with the Fluorescence polarization assay. We also thank the NIH Tetramer Core Facility (NIH Contract 75N93020D00005 and RRID: SCR_026557) for providing PE-labeled H-2k(b)-OVA_257-264_ and PE-labeled H-2D(b)-Adpgk (MC38) tetramers.

## Author contributions

Conceptualization: SSYL, SOL, DJ. Study design: DJ, SSYL, SOL. Antibody production: SOL, NL. Biophysical characterization, cellular experiments and in vivo experiments: DJ, XC, ZW. Microscopic imaging and image data analysis: XC, DJ, NR, YH. Writing-original draft: DJ, SSYL. Writing-review&editing: All authors

## Data and materials availability

All data are available in the main text or the supplementary materials.

