## Supplementary Data for "CD40 agonistic-monovalent streptavidin fusion antibody for targeted neoantigen peptide delivery and potent cancer vaccination"

Dahee Jung *et al.*

**This PDF file includes:**

Figs. S1 to S7
Table S1 to S3

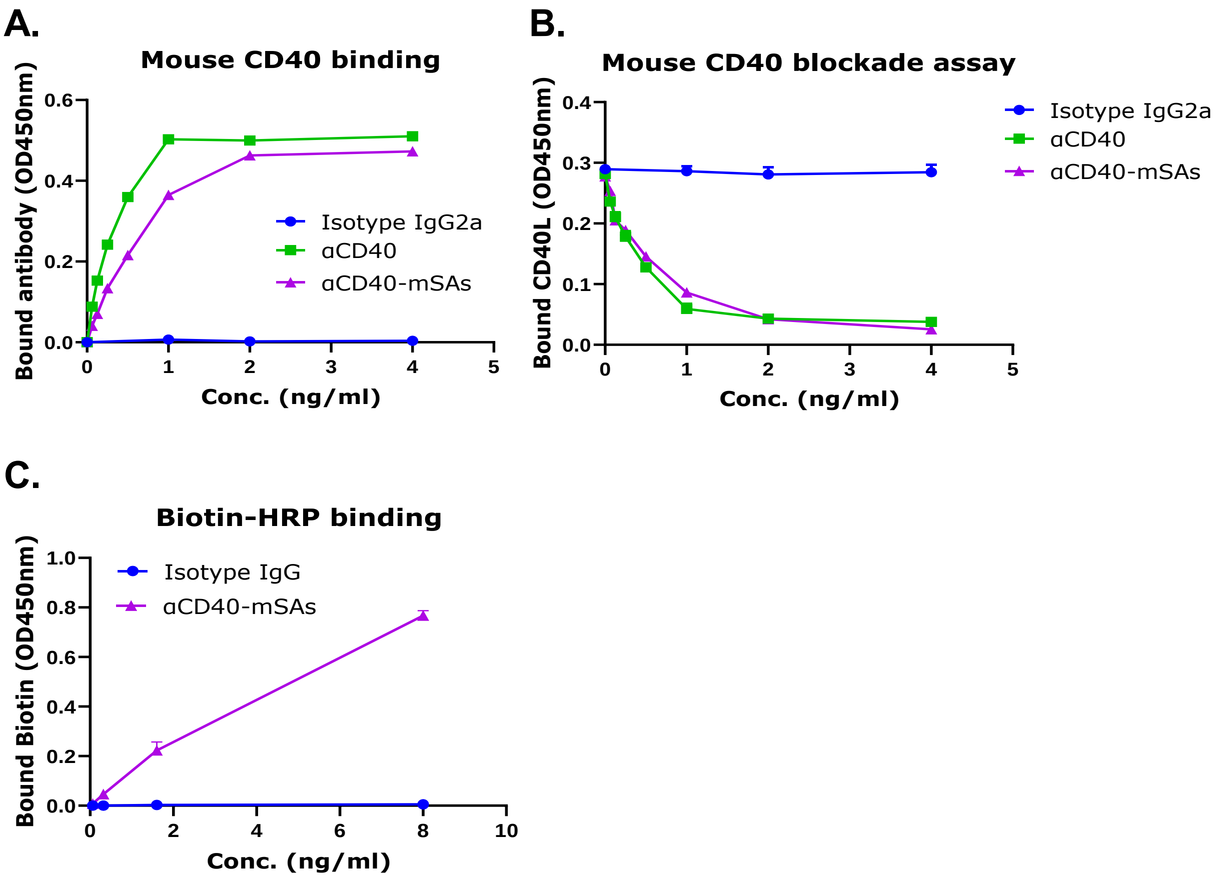


**Figure S1. ELISA assays evaluating the binding of recombinant αCD40 and αCD40-mSAs to mouse CD40 and biotin.** (A) Dose–response ELISA assessing the binding specificity of αCD40 and αCD40-mSAs to mouse CD40 (mCD40). The half-maximal binding concentrations (KD) for αCD40 and αCD40-mSAs were 0.3455 ng/mL and 0.5546 ng/mL, respectively. (B) Competitive binding assay (sandwich ELISA) further confirming the specific binding of αCD40 and αCD40-mSAs to mCD40. The bound mouse CD40L-Fc protein on mouse CD40 protein was quantified by measuring OD450. (C) The biotin binding ability of αCD40-mSAs was confirmed by measuring the HRP-conjugated biotin bound to αCD40-mSAs. Mouse IgG2a served as a negative control in all (A, B, and C)

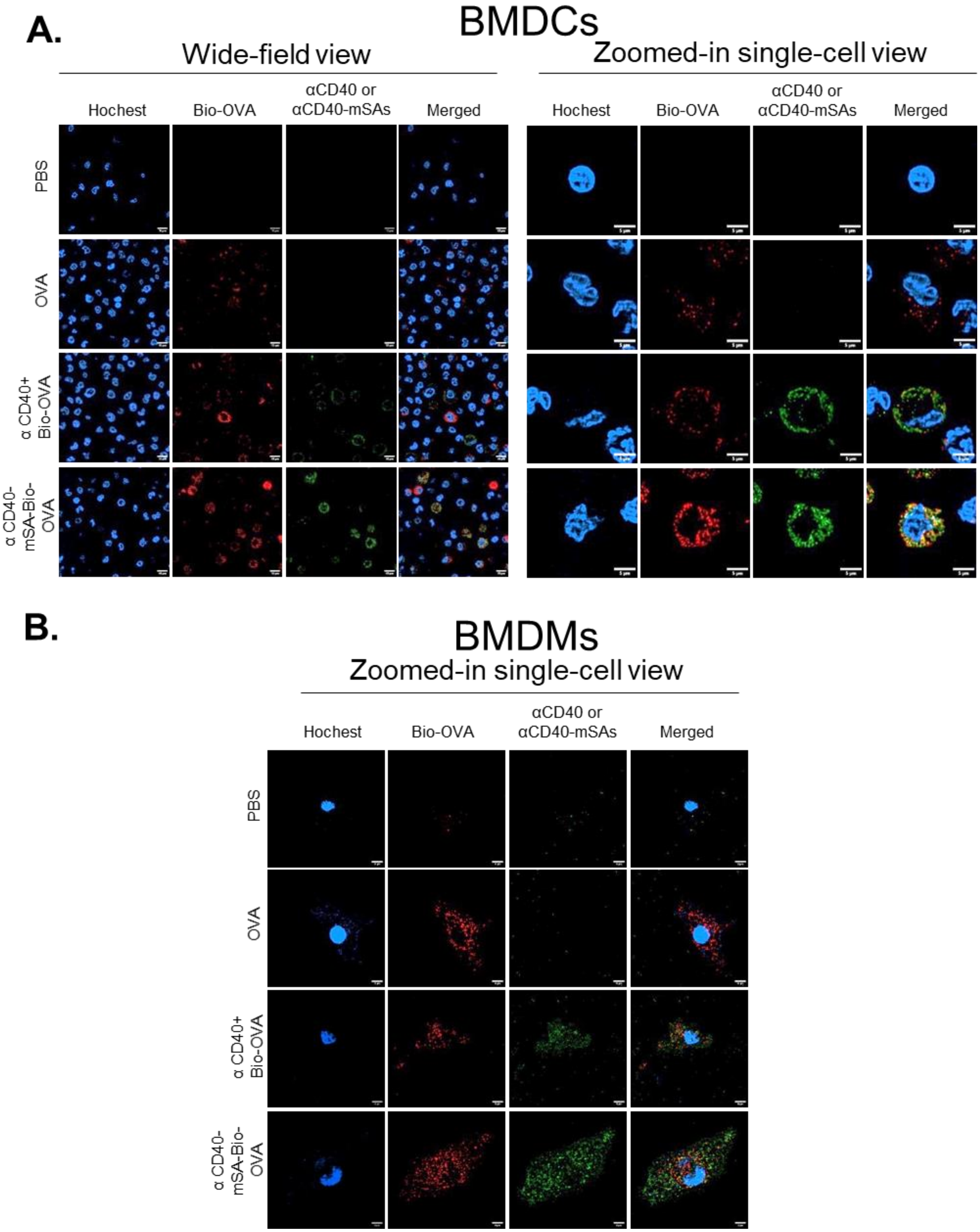


**Figure S2. Deep-SIM images** of (A) BMDCs and (B) BMDMs after 1-hour incubation with Bio-OVA-AF647 vaccine formulations containing either αCD40-DL550 or αCD40-mSAs-DL550. Scale bar: 15 μm (wide-field) 5 μm (Zoom-in single-cell).

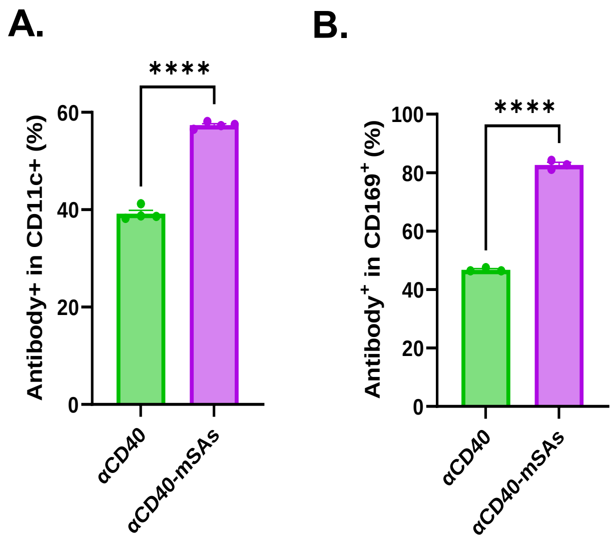


**Figure. S3. Flow cytometry analysis** of αCD40-DL550 and αCD40-mSAs-DL550 internalization into (A) BMDCs and (B) BMDMs after 1-hour incubation with Bio-OVA-AF647 vaccine formulations.

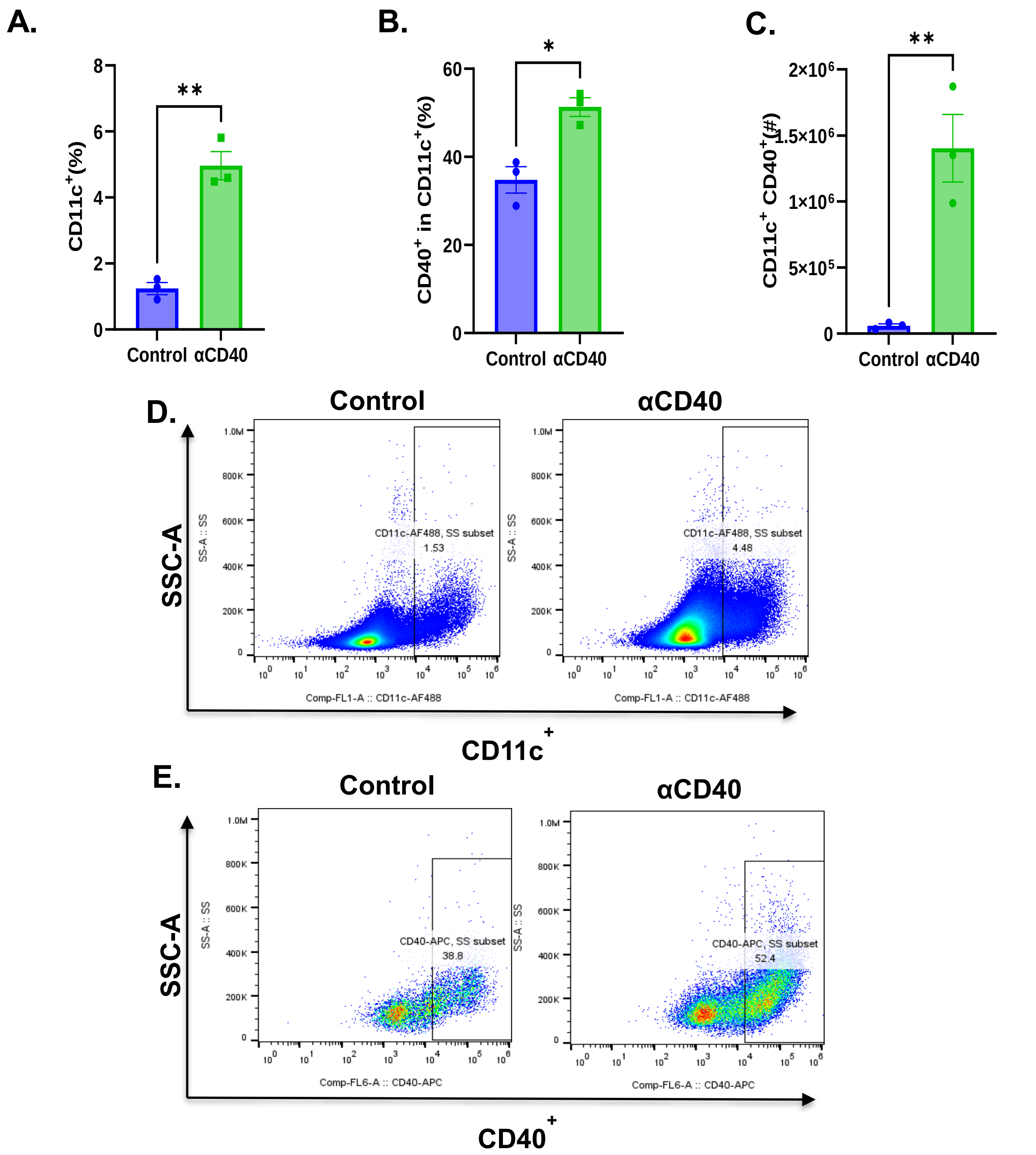


**Figure S4.** ***In vivo*** **adjuvant effect of αCD40 injection.** Mice (n = 3) received a single dose of αCD40 (100 µg in 30 µl) *via* footpad injection. Popliteal, inguinal, and axillary dLNs from both sides were collected 24 hours post-injection. (A) Percentage of CD11c$^{+}$ DCs among total cells in the collected dLNs. (B) Percentage of CD40$^{+}$ cells within the CD11c$^{+}$ DC population. (C) Absolute number of CD40$^{+}$ CD11c$^{+}$ DCs in the dLNs. (D, E) Representative flow cytometry plots showing CD11c$^{+}$ cells (D) and CD40$^{+}$ cells within the CD11c$^{+}$ DC population (E).

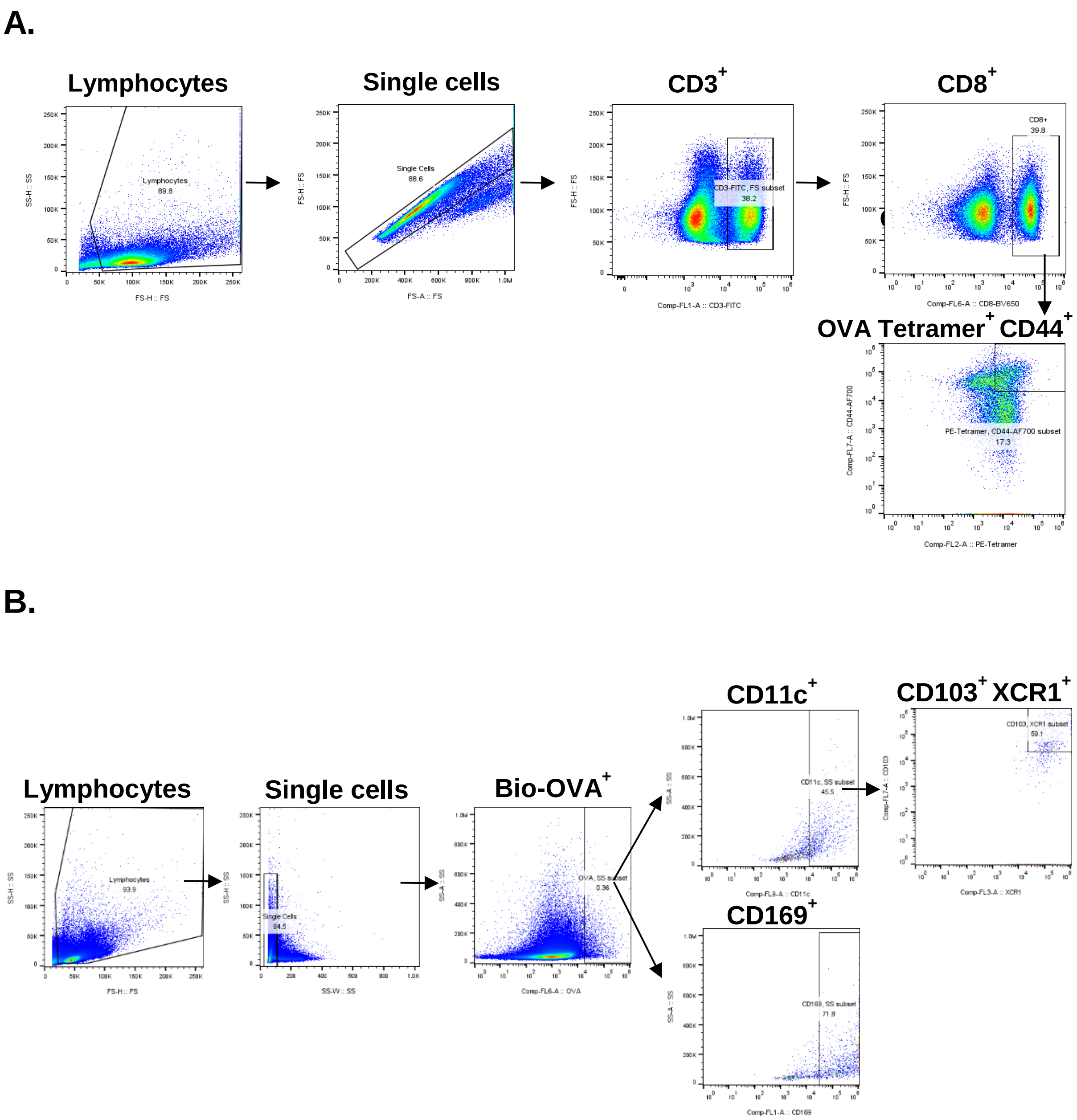


**Figure S5. Gating strategy for the study of (A) OVA specific T cells in dLNs and spleen from Figure 2 and (B) Distribution of Bio-OVA in dLNs from Figure 3.**

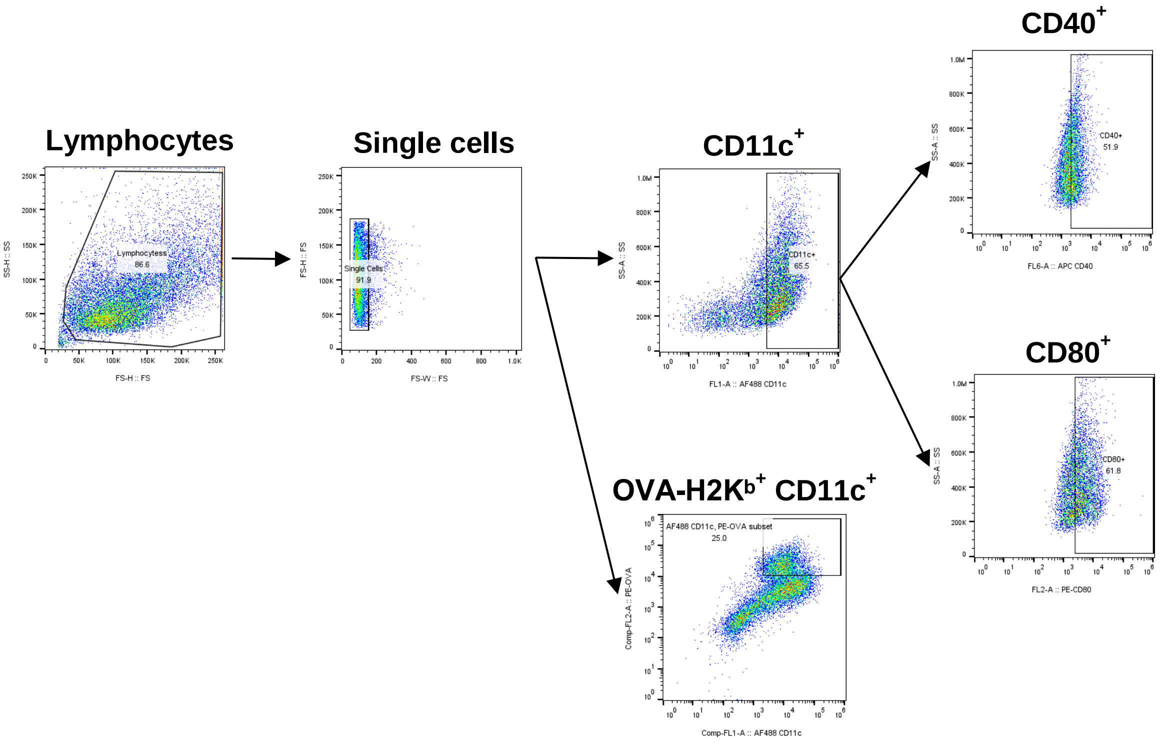


**Figure S6. Gating strategy for the study of BMDCs activation and antigen presentation from Figure 4.**

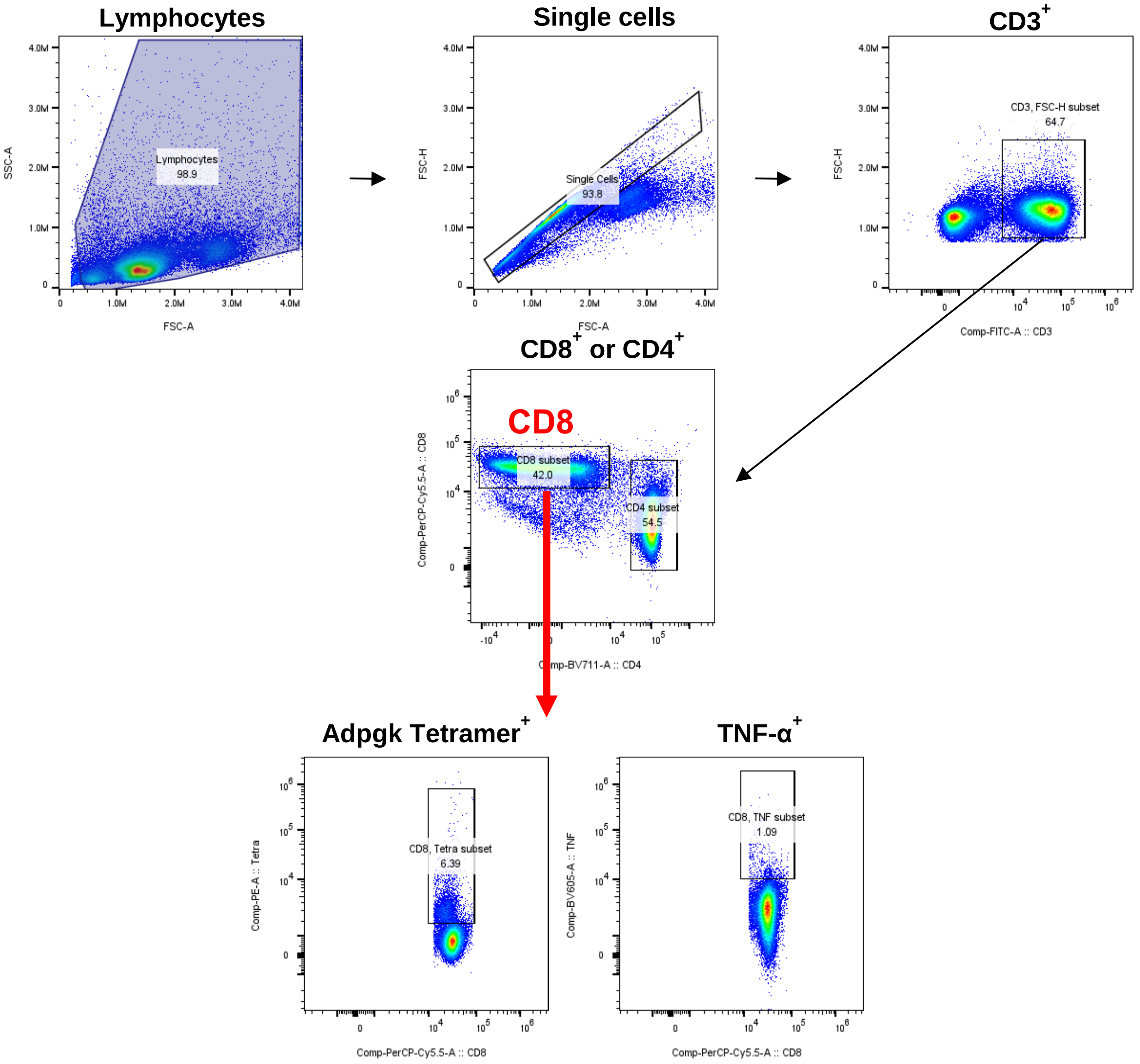


**Figure S7. Gating strategy for the study of T cell profiling from Figure 7.**

**Table S1. Neoantigen peptide information**

| Peptide name | Sequence | Molecular Weight |
|---|---|---|
| Bio-OVA$_{247–264}$-A5K (Bio-OVA) | Biotin-DEVSGLEQLESIINFEKLAAAAAK | 2772.6 |
| Bio-OVA$_{247–264}$-A5K-AF647 (Bio-OVA-AF647) | Biotin-DEVSGLEQLESIINFEKLAAAAA-Lys(N3)-Alexa647 | 3916.48 |
| Bio-MPEG-Adpgk-K5 (Bio-Adpgk) | Biotin-{mini PEG3}-HLELASMTNMELMSSIVHQKKKKK | 3241.87 |

**Table S2. List of antibodies and reagents used for flow cytometric assays.**

| Marker | Fluorophore | Clone | Dilution or amount | Catalog number | Vendor |
|---|---|---|---|---|---|
| CD11c | Alexa Fluor® 488 | N418 | 1:200 | 117311 | Biolegend |
| CD11c | BV421 | N418 | 0.5 µg | 117329 | Biolegend |
| CD169 | AF488 | 3D6.112 | 0.5 µg | 142419 | Biolegend |
| CD103 | Alexa Fluor® 700 | 2E7 | 1.25 µg | 121442 | Biolegend |
| XCR1 | PE/Dazzle594 | ZET | 0.3 µg | 148234 | Biolegend |
| CD40 | APC | 3/23 | 1.25:100 | 124612 | Biolegend |
| CD80 | PerCP/Cyanine5.5 | 16-10A1 | 2.5:100 | 104707 | Biolegend |
| OVA$_{257-264}$/H-2K$^{b}$ | PE | 25-D1.16 | 0.625:100 | 141603 | Biolegend |
| Live/Dead | Zombie violet | | 1:1000 | 423114 | Biolegend |
| CD3ε | FITC | 145-2C11 | 2:100 | 100203 | Biolegend |
| CD4 | BV711 | RM4-5 | 5:100 | 100550 | Biolegend |
| CD8α | PerCP/Cyanine5.5 | 53-6.7 | 4:100 | 100734 | Biolegend |
| CD44 | Alexa Fluor® 700 | IM7 | 0.5:100 | 103029 | Biolegend |
| CD62L | BV650 | MEL-14 | 2.5:100 | 104453 | Biolegend |
| TNF-α | BV605 | MP6-XT22 | 4:100 | 506329 | Biolegend |
| TruStain FcX™ (anti-mouse CD16/32) | | 93 | 2:100 | 101320 | Biolegend |
| H-2k(b)-OVA$_{257-264}$ | PE | Tetramer | 1:100 | | NIH tetramer core Facility |
| H-2D(b)- Adpgk (ASMTNMELM) | PE | Tetramer | 1:100 | | NIH tetramer core Facility |

| | | | | | |
|---|---|---|---|---|---|
| Cyto-Fast™ Fix/Perm Buffer Set | | | | 426803 | Biolegend |

**Table S3. List of antibodies and reagents used for microscopic imaging assays**

| **Marker** | **Fluorophore** | **Clone** | **Dilution** | **Catalog number** | **Vendor** |
|---|---|---|---|---|---|
| CD11c | Alexa Fluor® 488 | N418 | 0.2 mg/mL | 117311 | Biolegend |
| CD169 | Alexa Fluor® 488 | 3D6.112 | 0.02 mg/mL | 142419 | Biolegend |
| Cell nuclei | Hochest 34580 | | 0.1 mg/mL | H21486 | Invitrogen |
